# Polarizing SNPs without outgroup

**DOI:** 10.1101/2025.01.31.635919

**Authors:** Jinyang Liang, Julien Y. Dutheil

**Affiliations:** Department of Theoretical Biology, Max Planck Institute for Evolutionary Biology, August-Thienemann-Straße 2, 24306 Plön, GERMANY

**Keywords:** ancestral recombination graph, unfolded site-frequency spectrum, polarization, outgroup

## Abstract

Asserting which allele is ancestral or derived, known as polarization, is a prerequisite of many population and quantitative genetic methods. In particular, it allows inference of the unfolded site-frequency spectrum (uSFS). The most widely used approaches are based on outgroup data. However, for studies on species for which closely related outgroups are difficult to obtain, information on many sites of interest may be missed due to alignment problems. Here, we present PolarBEAR (Polarization By Estimation of the Ancestral recombination graph), a method that uses the local genealogies from the ancestral recombination graph (ARG) to infer ancestral states. We show that PolarBEAR can reach high accuracy in polarization and uSFS estimation using simulations under several scenarios. This accuracy, however, heavily depends on the ARG used as input. It is maximal when the true ARGs is used, but can be very low depending on the ARG reconstruction method employed. We also applied our method to human population data and compared it with the outgroup-based method est-sfs. Although our method could not infer the ancestral state with high confidence at certain positions, it obtained results for positions that est-sfs could not polarize due to missing outgroup data. The polarization results of the two methods were highly consistent at positions inferred by both methods. The two methods inferred similar uSFS, with PolarBEAR estimating slightly fewer high-frequency derived alleles. Furthermore, we demonstrate that PolarBEAR is robust to the mutation model used, while est-sfs exhibits a bias in the presence of heterogeneous base composition. PolarBEAR can complement outgroup-based methods, or replace them when no appropriate outgroup sequence is available.

## Introduction

The process of inferring the ancestral vs. derived state of alleles at polymorphic sites is called “polarization”. Polarized alleles are a required prior knowledge for many population genetics studies. Ancestral alleles are required as input to ARG inference methods such as Relate (Speidel et al. 2019) and tsinfer (Kelleher et al. 2019). This knowledge can also be used to detect adaptive evolution (Koenig et al. 2019), introgression (Fang et al. 2024), estimate allele age (Albers and McVean 2020; Pivirotto et al. 2024) and the strength of local selection (Stern et al. 2019). Furthermore, many methods have been developed to infer population genetic parameters from the distribution of the complete derived allele frequency spectrum, the so-called unfolded site frequency spectrum (uSFS), calculated from polarization results and containing signatures of demographic change (Keightley and Eyre-Walker 2007; Gutenkunst et al. 2009), gene flow (Marchi and Excoffier 2020) and selection (Zeng et al. 2006; Keightley and Eyre-Walker 2007; Schneider et al. 2011; Sendrowski and Bataillon 2024).

Ancestral or derived states cannot be directly observed from sequencing data. Current approaches consist in using the information from a homologous sequence of a closely related outgroup species. The simplest approaches equal the outgroup and the ancestral states (Voight et al. 2006; Schaefer et al. 2021), ignoring any evolutionary events since the common ancestor on the outgroup branch. Some early methods based on several outgroups and parsimony to infer ancestral states have been proposed (Sabeti et al. 2007; Langley et al. 2012), but other studies have pointed out their limits (Felsenstein 1981; Collins et al. 1994; Eyre-Walker 1998). These methods tend to reconstruct common nucleotides as ancestral states, resulting in an overestimation of the number of rare changes. Furthermore, mis-polarized variants can cause serious deviations in population genetics parameters estimation in downstream analyses and bias the conclusions (Hernandez et al. 2007; Glémin et al. 2015). Currently, the state-of-the-art method for ancestral state inference is est-sfs (Keightley et al. 2016; Keightley and Jackson 2018), based on maximum likelihood inference with up to three outgroups, which largely eliminates the bias compared to the parsimony method.

However, these outgroup-based methods all have a common problem: due to species divergence, outgroup data are missing for some variants; an effect even stronger when multiple outgroups are used. Hernandez et al. used chimpanzee and rhesus macaque as outgroups to demonstrate the impact of this problem on the analysis of human population data. Reliance on outgroups makes the situation more difficult in the cases when (1) it is difficult to obtain suitable outgroup data for the focal species, such as when the focal species is a relict or non-model species, and (2) it is more challenging to obtain homologous sequences with outgroups, such as in non-conserved or rearranged genome regions.

Here we develop an outgroup-free method, PolarBEAR (Polarization By Estimation of the Ancestral Recombination graph), based on ARGs and an empirical Bayesian approach to infer ancestral states and estimate uSFS. We assess the accuracy, power, and limits of this new method in comparison with classical approaches on simulated data and apply it to human population data from the 1000 Genomes Project as a test case.

## Results and discussion

### Inferring ancestral alleles using the ancestral recombination graph (ARG)

The ancestry of a sample of sequences is a complex entity represented by the ARG. It records two types of events: coalescence between lineages and crossing-overs. At any given position in the genome, the marginal genealogy is a rooted tree with branch lengths representing the ages of the ancestral genotypes. Given the local genealogy, it is possible to reconstruct the ancestral alleles at each coalescence node, including the root node. Model-based ancestral state reconstruction procedures have been developed for phylogenetics (Yang et al. 1995). These methods require a substitution model to compute transition probabilities along each tree branch. An empirical Bayesian approach is then used to compute the posterior probability of each possible ancestral state at each node with joint reconstruction (Yang et al. 1995).

Inferring the ancestral state at the most recent common ancestor of the sample may be challenging. When the history of the sample contains a single mutation, uncertainty happens when the mutation occurs on one of the branches at the root node (Figure 1A). The inferred ancestral state depends on whether the mutation occurs left or right of the root node, both scenarios being equally parsimonious. In terms of likelihood, the probability of each scenario only depends on the branch lengths and the chosen substitution model, resulting in very similar posterior probabilities. We refer to such scenarios as non-informative genealogies (with respect to ancestral allelic reconstruction), as opposed to informative genealogies when mutations occur on branches not connected to the root. When multiple mutations occur, more ambiguous scenarios can be observed.

**Figure 1.**
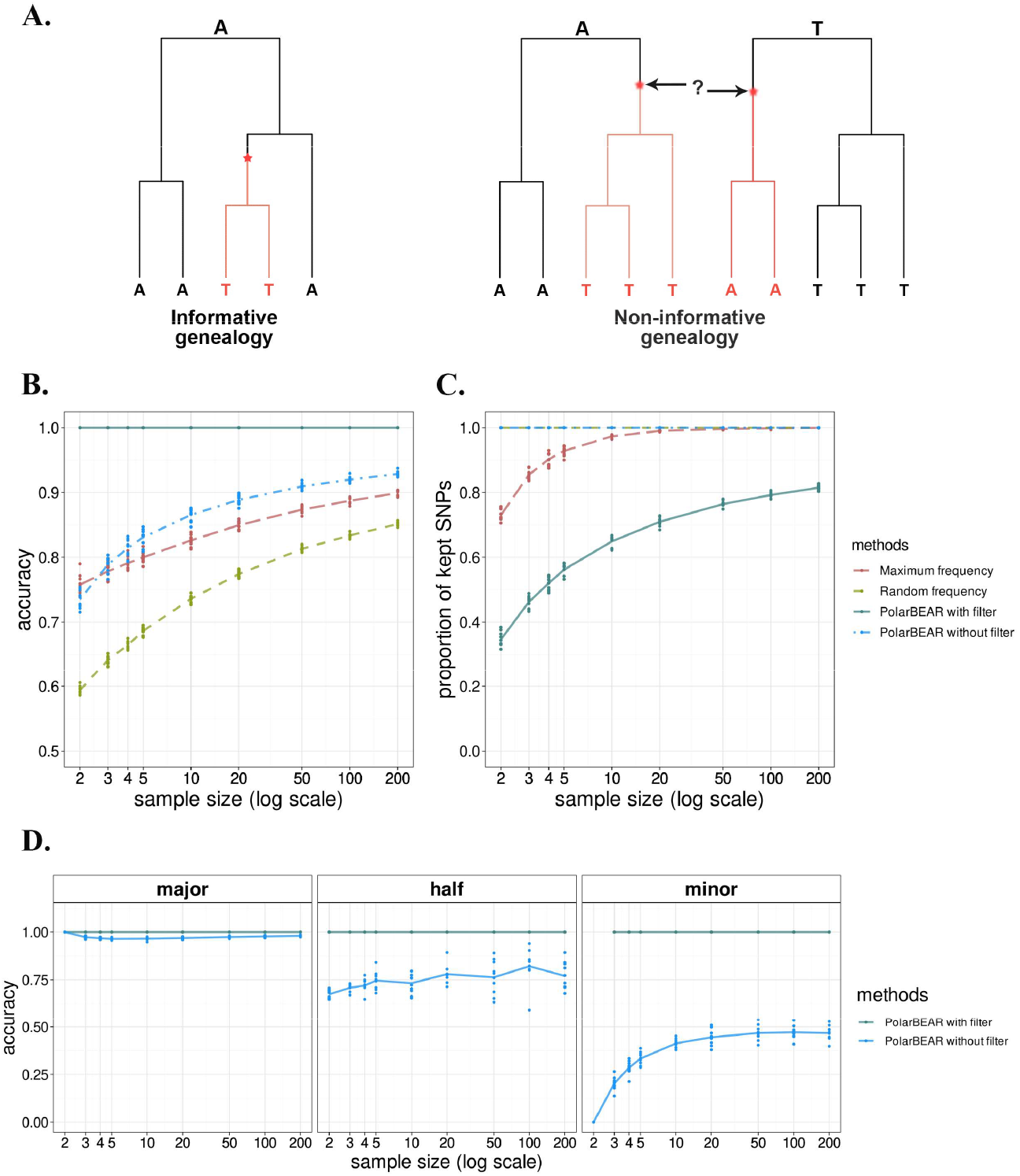
Polarizing performance of PolarBEAR with simulated ARG. A: examples of informative and non-informative ARG. The red stars and derived branches show the location of mutations. B and C: comparison of accuracy, that is, the frequency of correctly inferred ancestral alleles (B) and proportion of analyzed SNPs (C) between PolarBEAR with and without filtering of non-informative ARG, frequency-based and random methods (color) with different sample sizes. D: accuracy of PolarBEAR with and without filtering of non-informative genealogies among the SNPs classified by their ancestral alleles as major, equal, and minor for different sample sizes. The points show 10 replicates in each group, and the lines represent their means.

### Accuracy of allele polarization when the ARG is known

We first assessed the accuracy of the ancestral reconstruction in the ideal scenario where the true ARG is known and there is only one mutation per site (Figure 1B), using simulations (see Methods). We further demonstrate the impact of filtering out the non-informative genealogies (Figure 1B). The proportion of accurately inferred ancestral states (hereby referred to as “accuracy”) increases with the sample size. The accuracy of PolarBEAR is generally higher than considering the Major allele as the ancestral one (hereby referred to as the “Maximum frequency” method) or sampling according to the allelic frequency (hereby referred to as the “Random frequency” method). We note that for sample sizes above 20 diploid individuals, the average accuracy of the two latter methods is virtually identical on average. Using PolarBEAR while filtering out the cases of non-informative genealogies leads to an accuracy of 100% for all sample sizes, showing that when the ARG is known, the non-informative topologies account for the total of the erroneous inference. The proportion of positions with non-informative topologies is very high for small sample sizes (65% of the topologies for two diploids). It decreases as the sample size increases (20% of positions for 200 diploids, Figure 1C).

We then compare the inference with the frequency methods in more detail. By construction, the maximum frequency method will have an accuracy of 100% when the true ancestral allele is the major one and 0% when the true ancestral allele is the minor one. Using the true genealogy, PolarBEAR recovers the correct ancestral state with 100% accuracy when the ancestral allele is the major one, but the accuracy drops to 75% when the two alleles are in equal frequencies and to below 50% when the ancestral allele is the minor one (Figure 1D). In the latter case, the accuracy is even lower when the sample size is smaller than ten diploid individuals. When filtering out non-informative genealogies, the accuracy becomes 100% in all three cases, demonstrating that erroneous inferences stem from non-informative genealogies, which are more frequent when the ancestral allele is the minor one. The proportion of these genealogies increases with smaller sample sizes and reaches 100% for two diploids (if the minor allele is the ancestral one, it is in frequency ¼, and the corresponding individual is necessarily on a branch connected to the root). In subsequent results, we filter the non-informative genealogies when polarizing SNPs with the PolarBEAR method.

Finally, we assess the impact of multiple mutations on the ancestral state reconstruction (Figure S1). To obtain enough sites with multiple mutation events, simulations with a higher mutation rate of 1e-7 bp^-1^ were employed (see Methods). PolarBEAR infers the ancestral allele with an accuracy of more than 90% in samples of more than ten individuals when more than two alleles are present (Figure S1A). This means that the assumption of an infinite site model is not necessary when the ARG is accurate. When we further exclude the genealogies with homoplasy (two alleles but multiple mutations), the accuracy reaches 100% (Figure S1A and C), indicating that the same mutation occurring in different branches is the main reason for incorrect inference. The effect is stronger when the true ancestral allele is the minor one. Although the accuracy increases with sample size regardless of the ancestral allele frequency, it does not exceed 75% at a sample size of 20 when the ancestral allele is the non-major one. In comparison, it is close to 100% when the ancestral is the major one for all the sample sizes (Figure S1C).

### Accuracy when the ARG is inferred from the data

We next examine how the method performs in practical cases when the ARG is unknown and must first be inferred from the data. It should be emphasized that due to the principle of ARG, the data must be phased. ARGs can be directly obtained from methods such as tsinfer and RENT+ (Mirzaei and Wu 2017), or reconstructed from the set of times to the most recent common ancestor (TMRCA) of every pair of genomes estimated using sequentially Markovian coalescent (SMC)-based methods (e.g., PSMC, Gamma-SMC) (Li and Durbin 2011; Schweiger and Durbin 2023). For the latter, the TMRCA of all pairs of haplotypes in the sample at segregating sites are combined into distance matrices, which are subsequently used to reconstruct the rooted local genealogies with the unweighted pair group method with arithmetic mean (UPGMA) algorithm. As tsinfer only reconstructs the tree topology, tsdate (Wohns et al. 2022) is used next to determine branch lengths (hereby referred to as “tsinfer+tsdate”). We compare different reconstruction methods and assess their impact on the ancestral allele reconstruction. It should be noted that tsinfer requires ancestral alleles as input. In our experiment, we used major alleles as the ancestral ones to see whether it could obtain accurate ARGs and polarized results under this premise. For PSMC, the maximum sample size used was ten diploids due to computational resource limitations and 100 for the other methods.

Unlike the true ARGs, the inferred ARGs contain uncertainty and topological errors. The uncertainty is reflected in the fact that some topologies contain polytomies, especially in the results of methods such as RENT+ and tsinfer+tsdate that directly infer ARGs for the full samples. When mutations occur at polytomies, we cannot rule out that the underlying true genealogies are non-informative. However, topological errors may not affect polarization, for instance, when they are under a common ancestor node without any mutations below it. In other cases, they may result in an unreasonably high number of inferred mutation events. We reconstructed the positions of mutations along the inferred genealogies and filtered out cases where mutations occurred under a polytomy or when multiple mutations were inferred. The first filtering scheme mainly affects RENT+ and tsinfer+tsdate, as mentioned above, while the latter is useful for PSMC and gamma-SMC, both of which have more noticeable effects when the true ancestral allele is the minor one or half-half (Figure S2). When the sample size is two, all the genealogies in ARGs obtained by tsinfer are polytomies at the root node, so when the first filter is implemented, no analyzable SNPs are kept (Figure S2 and Figure 2B).

**Figure 2.**
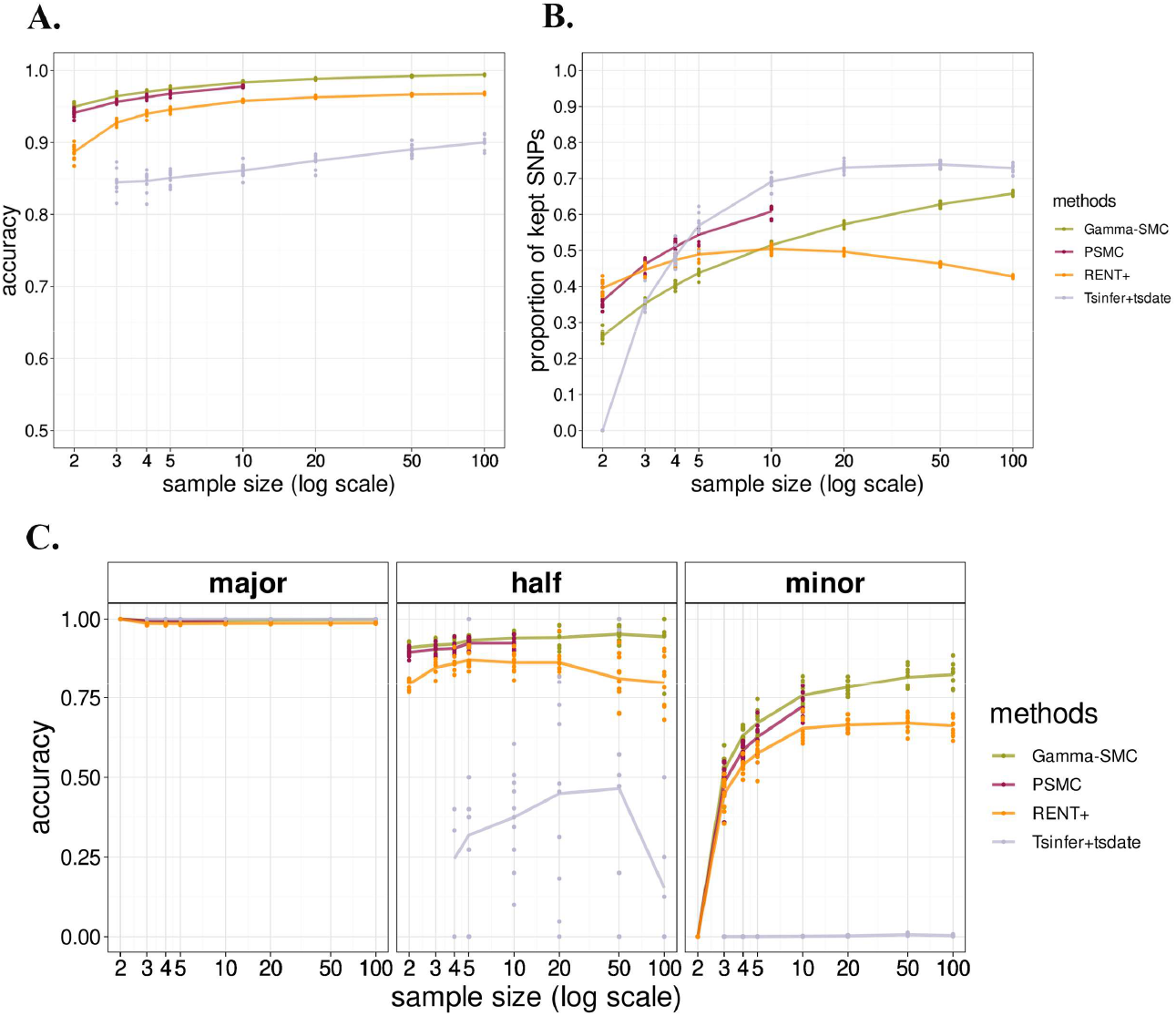
Polarizing performance of PolarBEAR with inferred ARGs. ARGs inference methods include gamma-SMC, PSMC, RENT+, and tsinfer+tsdate, shown in different colors after applying filters. Comparison of total accuracy (A), proportion of kept SNPs (B) and accuracy of SNPs classified according to their ancestral alleles’ frequencies (major, equal, and minor, C.) for different sample sizes. The points show 10 replicates in each group, and the lines represent their means. Missing values are reported when no sites are left after filtering.

We compared the accuracy of these methods after implementing all filters and found that the accuracy of gamma-SMC, PSMC and RENT+ can reach more than 95% when the sample sizes are larger than ten individuals, while tsinfer+tsdate only reaches 90% accuracy when the sample size is 100. (Figure 2A). The accuracy of all methods increases with the sample size, among which gamma-SMC and PSMC have the highest accuracy (≥97% for a sample size of ten, with gamma-SMC being slightly more accurate). When the true ancestral allele is the major one, all methods approach 100% accuracy (Figure 2C). When the frequencies of ancestral and derived alleles are equal, the variation in accuracy between different replicates of all methods becomes larger with increasing sample size because such sites become rare, especially after filtering is implemented. The advantage of gamma-SMC is more pronounced when the true ancestral allele is the minor one, with an accuracy of 75% when the sample size is larger than ten. When the ancestral allele frequency is in the minority and half, the ranking of the accuracy of different methods is consistent with that of the total accuracy. On the other hand, PSMC has an advantage in the proportion of kept SNPs, at least in the case of small sample sizes (<= 10), as it systematically retains 10% more sites than the gamma-SMC method (Figure 2B). The poor performance of tsinfer+tsdate may be due to its reliance on the input ancestral allele. As we used the major allele as input ancestral state, it is incorrect for positions where the true ancestral allele is the minor one. This may explain why it has 100% accuracy when the ancestral alleles are the major ones but close to zero when the ancestral alleles are the minor ones (Figure 2C). We note that using the resulting ARG to re-infer the ancestral alleles did not lead to any further improvement (Figure S3A and B). In contrast, when using the true ancestral alleles as input, the accuracy was above 90%, even when the ancestral allele was the minor one (Figure S3C). Moreover, when the sample size is two, unlike the case mentioned above where major alleles are used as input, inferred genealogies no longer display polytomies at the root nodes and the proportion of analyzable sites increases from 0 to 36% (Figure S3D).

To test the performance of polarization under different demographic scenarios, we simulated a logarithmic change of about 50 times within 150,000 generations for both the population size increase and decrease scenarios. When the population size decreases, the overall accuracy of the methods does not change significantly. However, PSMC, previously close to gamma-SMC, now has the highest accuracy (Figure S4A). When the ancestral allele is the minor one, the accuracy of gamma-SMC decreases more obviously, and that of PSMC exceeds it by more than 5% (Figure S4C). Furthermore, the proportion of SNPs kept after filtering of PSMC and RENT+ decreased with respect to the scenario of constant population size, while it did not change significantly for gamma-SMC (Figure 2B and S4B). In the scenario where the population size increases, the overall accuracy of the methods is slightly higher than in the constant population size scenario (Figure 2A and S5A). However, this difference vanishes when classifying SNPs according to the frequency level of their ancestral alleles, suggesting that the average gain in accuracy is due to a higher proportion of SNPs where the ancestral allele is the major one in the population expansion scenario (Figure 2C and S5C). Furthermore, as opposed to when population size declines, the proportion of SNPs kept after filtering was higher for all methods in the growth scenario, with that of PSMC showing the largest increase (more than 10% increase, Figure 2B and S5B).

### Impact of the recombination rate on the ancestral allele reconstruction

The accuracy of ARG inference methods is known to be affected by the ratio of recombination rate to mutation rate. For a given mutation rate, the higher the recombination rate, the more recombination events will be unseen in the data. To understand how the recombination rate affects ARG inference and its impact on the polarization results, we classified the SNPs according to the local recombination rate (see Methods). The accuracy ranking of the four methods was consistent with previous results regardless of the recombination rate; and all methods had higher accuracy in regions with the lowest recombination rates, with tsinfer+tsdate having the smallest difference at 1% to 2% in accuracy between the two recombination rate classifications for all sample sizes (Figure S6A). The other three methods had greater differences in accuracy between low and high recombination rate regions, and these differences decreased with the sample size. For example, when the sample size was ten, the difference in accuracy between the two recombination rates of gamma-SMC was more than 12%, that of PSMC was close to 14%, and it was up to 15% for RENT+; when the sample size was 100, this difference was only 1% for gamma-SMC and RENT+. The sample size effect of the accuracy difference mainly comes from the improvement of accuracy with the increase of sample size at high recombination rates, while the accuracy under low recombination rate remains at a high level under most sample sizes: the accuracy of gamma-SMC and PSMC is above 98% in all sample sizes, and that of RENT+ is above 95% when the sample size was greater than two.

The impact of the recombination rate is also stronger when the ancestral allele is the minor one: for a sample size of ten, the difference in accuracy between the two recombination rates of RENT+ reaches 27%, and for gamma-SMC and PSMC it can even exceed 42% and 47%, respectively. (Figure S6C).

Recombination rate does not only impact the reconstruction accuracy but also the number of SNPs for which reconstruction can be performed. Some of the incorrectly reconstructed topologies will lead to a high number of mutations and will be filtered out, leading to a lower proportion of kept SNPs. The proportion of analyzable SNPs by gamma-SMC and PSMC increases with the sample size for both recombination rate scenarios but increases less when the recombination rate is high (Figure S6B).

Based on the above results, reconstructing the marginal genealogies using UPGMA on the matrix of pairwise TMRCAs inferred by gamma-SMC offers the best compromise in accuracy, running speed and memory usage. It notably permits the analysis of large sample sizes with good accuracy. When the sample size is not more than 10, PSMC is also a worthy alternative, given its accuracy close to that of gamma-SMC and a 10% higher proportion of polarizable SNPs.

### Estimating the unfolded site frequency spectrum

In this section, we present a scheme for estimating the unfolded site frequency spectrum (uSFS) using polarization posterior probabilities. We compared three alternative methods to infer the uSFS and used quantile-quantile (QQ) plots to compare the results with the true uSFS (Figure 3). Two of the three alternative methods directly use the polarization results mentioned in the previous sections: with several filtering (hereby referred to as the maximum posterior probability with filtering, “MAX PP + filtering” method); and without filtering of non-informative genealogies (hereby referred to as the maximum posterior probability, “MAX PP” method). The uSFS inferred with the MAX PP + filtering method significantly deviates from the true uSFS, even more than the uSFS inferred without filtering. This is because the number of non-informative genealogies depends on the derived allele frequencies, resulting in a bias when filtering genealogies (Figure S7).

**Figure 3.**
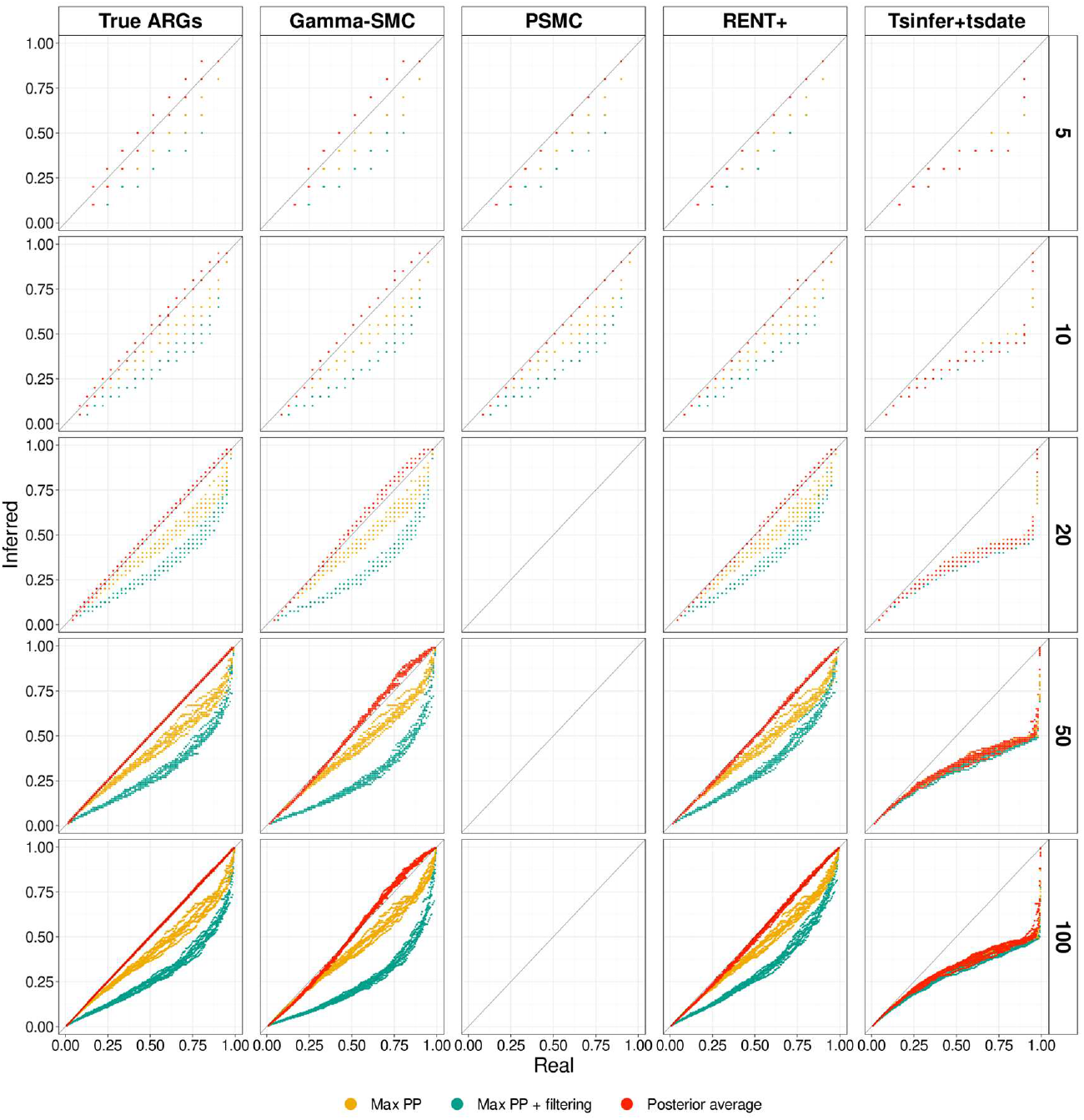
uSFS inference with different ARG reconstruction methods under a constant demography scenario. The panels show the comparisons of different uSFS computation methods with distinct colors, as quantile-quantile plots with the true uSFS, for different ARG reconstruction methods (columns) and sample sizes (rows).

We introduce a third method, hereby referred to as the “Posterior average” method. We include the uncertainty of ancestral allele reconstruction when computing the uSFS to account for possible polarization errors. From the results in the previous sections, a source of uncertainty is non-informative genealogies. We show that the relative branch lengths at the root node of these topologies contain information about the ancestral states, a signal captured by the posterior probabilities (Supp. Text 1). Using posterior probabilities as proxies for the polarization error rate, we can compute the expectation of the true proportion of sites in each derived allele frequency (DAF) category. We consider a sample of *n* haplotypes, bi-allelic polymorphic sites in DAF category *D*_*x*_ with frequency *x*/*n* have *x* copies of the derived allele and *n* − *x* copies of the ancestral allele. The expectation *E*[*D*_*x*_] of the true number of sites in category *D*_*x*_ is then computed as the sum of probabilities of each SNP *i* to be in the category. For instance, a SNP with two A alleles and eight T alleles is in category *D*_2_ if T is the ancestral allele and in category *D*_8_ if A is the ancestral allele. The probability that the SNP belongs to category *D*_2_, noted *P*(*D*_2_), is equal to the posterior probability that T is the ancestral state at this position. Similarly, *P*(*D*_8_), is equal to the posterior probability that A is the ancestral state at this position. Note that all other probabilities, *P*(*D*_l,3,4,5,6,7,9_) are null for this site.

We then have:

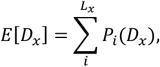

where *L*_*x*_ denotes the number of sites with allele frequencies (*x*/*n*, (*n* − *x*)/*n*), in whichever order.

Using the posterior-averaged uSFS with the true ARG leads to estimates that are only slightly biased in the high DAF categories, although with large variance for small sample sizes (Figure 3). This confirms that the branch length information in ARGs is included in the ancestral states posterior probabilities and can be used to provide an effective estimate for the uSFS.

We then examine the performance of four ARG inference methods (Figure 3). The genealogies with more than two mutations were filtered out because they were more likely to imply erroneous topologies and lead to incorrect posterior probabilities. We note *L*′_*x*_. the number of SNPs with allele frequencies (*x*/*n*, (*n* − *x*)/*n*) that are kept after filtering. We then obtain the expectation of the true number of the SNPs in category *D*_*x*_ with

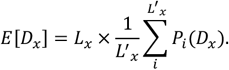

The uSFS count from ARGs inferred by gamma-SMC, PSMC and RENT+ closely followed the QQ-plot diagonal, but that of tsinfer+tsdate deviated significantly, possibly because of the incorrect input of major alleles as ancestral states. Among the first three methods, PSMC and RENT+ perform better, while the posterior average of gamma-SMC is slightly overcorrected.

Since the shape of the uSFS is strongly impacted by demography, we assessed the uSFS estimation accuracy under the same simulations of population expansion and contraction scenarios as in section *accuracy when the ARG is inferred from the data*. Under the scenario of decreasing population size, we observed an increased variance in the uSFS estimations, possibly due to the reduction in the number of SNPs. However, the inferred uSFS was very close to the true one, the RENT+ method leading to the best inference in this scenario, in particular when the sample size was large (Figure S8). In the scenario of increasing population size, the uSFS obtained using the ARG inferred from PSMC was close to the true one but the uSFS obtained from the ARGs inferred from RENT+ and gamma-SMC were biased (Figure S9). Among the three methods, gamma-SMC shows the largest departure from the true uSFS, possibly because it assumes a demographic model with constant population size in its flow field construction (Schweiger and Durbin 2023).

We note that, when using the true ARG, the uSFS is recovered with high accuracy by the posterior average method in all three demographic scenarios, suggesting that non-constant population sizes affect the ARG reconstruction itself rather than the polarization per se. Two properties of the inferred ARGs directly affect estimates of the uSFS: the number of inferred non-informative genealogies and the accuracy of branch lengths. While the posterior probabilities of the informative genealogies are usually close to 1 and will have little impact on the posterior averaging correction, non-informative genealogies have probabilities closer to 0.5, reflecting the uncertainty in the ancestral state reconstruction and impacting the uSFS estimation. While accounting for true non-informative genealogies enables unbiased uSFS inference, incorrect reconstruction of non-informative genealogies could lead to an over-correction, especially when the number of SNPs in the two complementary categories *D*_*x*_ and *D*_*n*-*x*_ is very different. Furthermore, even when non-informative genealogies are correctly recovered, underestimating the differences in branch lengths at the root node will push the posterior probability closer to 0.5, leading to a similar effect as overestimating the number of non-informative genealogies. To understand the source of the bias in uSFS estimates caused by the results of different ARG inference methods, we examined the difference between the non-informative genealogies proportion from inference and the true of non-informative genealogies (Figure S7). Among the three demographic scenarios, the proportion of non-informative genealogies inferred from PSMC is closest to the true one. The proportion of non-informative genealogies estimated by gamma-SMC is overestimated to different levels under the three demographic simulations: It is closest to the true one in the scenario of population contraction and greatly overestimated in the scenario of population expansion. In both cases, the overestimation is present in the most abundant low DAF categories. In contrast to gamma-SMC, RENT+ underestimates the proportion of non-informative genealogies under different demographic scenarios, an effect that increases with the sample size. However, in the scenario where the population size increases, the uSFS estimated using the results of RENT+ also shows a small amount of overcorrection compared to the MAX PP and MAX PP + filtering methods, which may be due to the differences in branch lengths at the root node being underestimated.

Since QQ-plots focus on global distribution similarity, and the number of sites in the highest DAF categories is small, the bias does not appear with such representation. We compute the relative error, calculated as the difference between the inferred frequency and the true frequency, divided by the true frequency, to evaluate the performance of PolarBEAR with PSMC, the best performing method mentioned above (Figure S10). In the scenario of decreasing population size, when the sample size was ten, the number of sites in each DAF categories was too small so the variation was large, making it difficult to determine whether it is biased, but when the sample size was five, the bias was the smallest among the three demographic scenarios; in the scenarios of constant and increasing population size, it showed an underestimation in the high DAF categories, which was most obvious in the last element, respectively at 0.13 and 0.16 when sample size was ten. However, these biases were absent when the true ARGs were used, indicating that they are caused by the inaccuracy of ARG inference by PSMC.

In outgroup-based uSFS estimation methods, misspecification of the substitution rate model will seriously affect the results (Keightley et al. 2016). It is worth mentioning that in our ARG-based uSFS estimation, although we assume the Jukes-Cantor model, the uSFS will not be biased when the ARG is accurate and the substitution rates between the two alleles are symmetric. Because in this case, in a non-informative genealogy, the rate difference of the mutation types can be scaled in the mutation rate *θ*, which is equal on both branches of the root node, and the posterior probability depends only on the ratio of the two branch lengths (Supp. Text 1). While the posterior probabilities of the informative genealogies are close to one and the impact can be ignored. We verified this using the simulations with Kimura’s two-parameter model (Kimura 1980) with the transition/transversion rate ratio *κ* = 5 (Figure S11). As expected, when the ARGs were known, the uSFS estimated by PolarBEAR was almost unbiased. When the ARGs were inferred from PSMC, the estimated uSFS was underestimated in the high DAF category, which is similar to the previous results and is probably due to inaccurate ARG reconstruction, but the relative error of uSFS did not differ among different mutation types.

The results in this section suggest that for the estimation of the uSFS based on the ARG, due to the uncertainty of the ancestral state at a specific site, directly using the most likely ancestral state will lead to a severe overestimation in low DAF categories and underestimation in high DAF categories, while combining with posterior averaging can well recover the uSFS. Additionally, the uSFS estimation using PSMC for ARG reconstruction performs best despite slightly underestimating the high DAF categories, followed by RENT+, while gamma-SMC leads to a relatively large bias.

### Performance on real data

We compare PolarBEAR with the outgroup-based method est-sfs on a benchmark data set. We used the human African population Mende, Sierra Leone (MSL) from the 1000 Genomes Project, containing 87 diploid individuals, restricting the analysis to chromosome 1. Orangutan and macaque were selected as outgroups for using with est-sfs. Sites with missing data in the outgroups and non-biallelic sites were discarded (Keightley & Jackson, 2018), and the R6 model, allowing six symmetrical rates, was used to estimate the substitution rates. We used PolarBEAR with all SNPs that pass all quality criteria (see Methods). ARGs were inferred using gamma-SMC and sites with non-informative genealogies or an inferred number of mutations greater than the number of observed alleles were filtered out.

We first examined the proportion of SNPs that were polarizable by each method. Then, among the SNPs polarizable by both methods, we focused on the proportion where the polarization results agreed to evaluate the consistency between the two methods. Est-sfs could analyze 86.4% of the input SNPs, while PolarBEAR could analyze a proportion of 60.3% of them. 52.2% of the SNPs could be analyzed by both methods, and 5.5% could not be analyzed by either PolarBEAR or est-sfs (Figure 4A). The two methods returned the same ancestral allele in 96.0% of the SNPs that could be analyzed by both methods. Using PolarBEAR in conjunction with est-sfs brings the proportion of polarizable SNPs from 86.4% to 94.5%.

**Figure 4.**
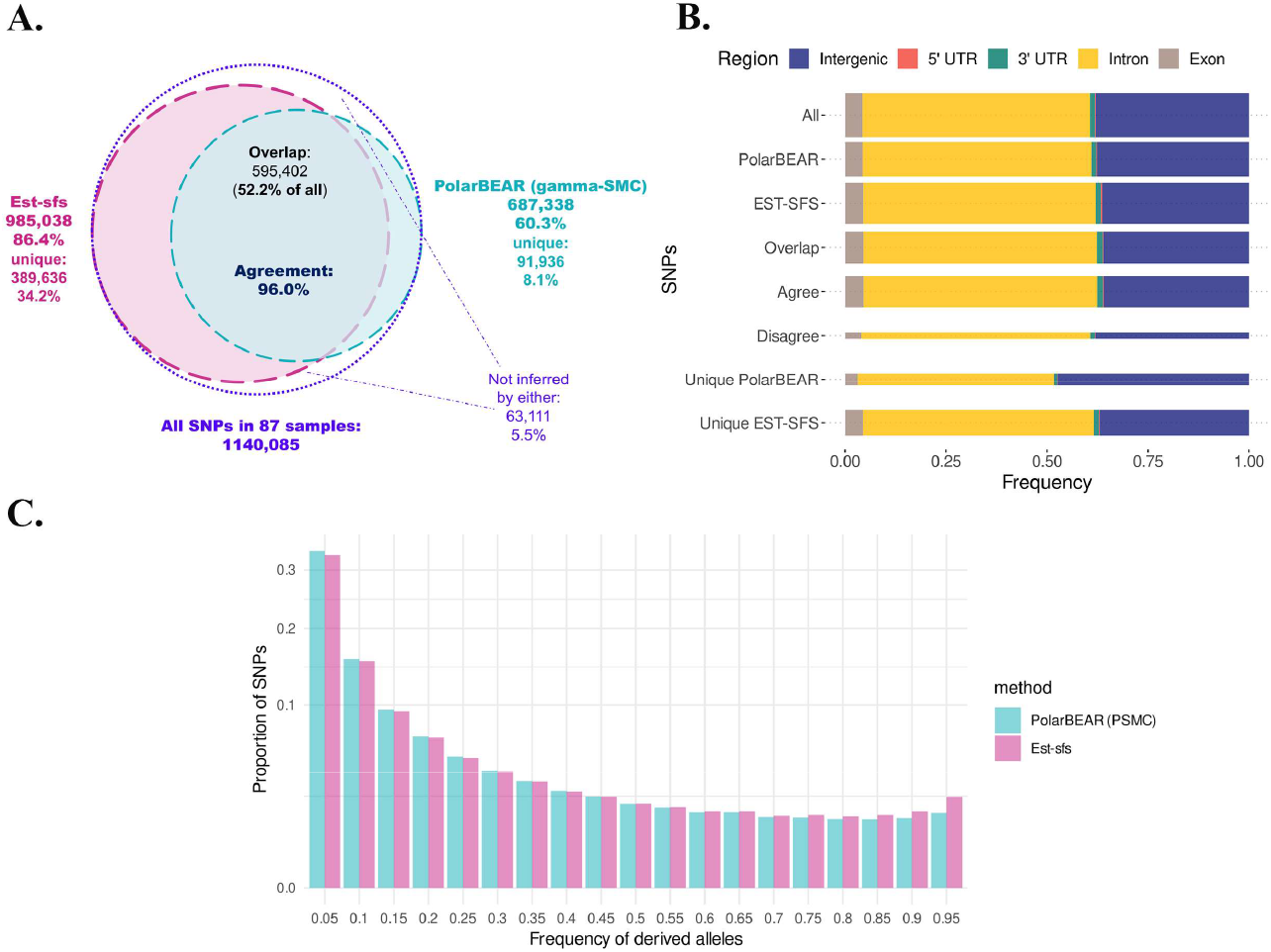
Comparison of Est-sfs and PolarBEAR polarization of SNPs of human chromosome 1. A: numbers and proportions of SNPs that can be analyzed by PolarBEAR, est-sfs and their overlap, as well as the agreement in the overlap between the two methods. B: Comparison of SNPs frequencies and methods agreement, in exon, intron, 5’ UTR, 3’UTR and intergenic regions. The widths of the boxes are proportional to the square roots of the numbers of SNPs in the groups. C: uSFS estimated by est-sfs (pink) and PolarBEAR with PSMC (blue). The y-axis (SNP proportion) is scaled using a square root transformation.

To examine the putative effect of selection on the performance of the two methods, we compared the SNPs in different regions expected to have different levels of selective pressure (Figure 4B). SNPs were classified into exons, 5’ UTRs, 3’ UTRs, introns, and intergenic regions. If selection affects the inference of PolarBEAR more than the reference method est-sfs, we expect that the distribution of SNPs for which the two methods provide consistent results will differ from the global distribution of SNPs analyzable by both methods. In particular, such consistently polarized SNPs should be under-represented in exonic regions, where selection is the strongest. First, while significantly different, the distribution of SNPs polarizable by PolarBEAR and est-sfs are very close to the global genome distribution (Figure 4B, chi2 test, P value = 6.647e-05 for PolarBEAR and 2.2e-16 for est-sfs). We note that the SNP distribution of est-sfs differs from the genome distribution to a larger extent than that of PolarBEAR. This difference is mostly imputable to a lower proportion of analyzable SNPs in intergenic regions. However, the frequency of agreement does not differ between genomic regions (chi2 test, distribution of SNPs where the two methods give a consistent result compared to the distribution of SNPs analyzable by both methods, P value = 0.8742). As intergenic regions evolve faster than coding regions, a comparatively higher fraction of positions cannot be aligned with the outgroup or may display alignment errors. As a result, we conclude that selection is not a major confounding factor during polarization using the ARG.

We inferred the uSFS with PolarBEAR using the ARG obtained with PSMC, as it was shown to provide the best performance. Sites that were inferred to have more than two mutations were excluded. We randomly selected ten individuals from the MSL population. In the middle part of the uSFS (DAF 0.3-0.7), PolarBEAR and est-sfs showed a strong agreement, but PolarBEAR inferred higher frequencies for low-frequency derived allele categories; whereas est-sfs inferred an increase in the frequencies of high-frequency derived variants (Figure 4C). Similar to the relative error calculated above, the uSFS from est-sfs was used as the “standard” to show the relative difference between PolarBEAR and est-sfs (Figure S12, black). Values of relative difference decreased with DAF and were negative in high DAF categories. Although part of it can be explained by the underestimation of PolarBEAR with PSMC, in the simulations with three different demographic scenarios and same sample size ten, none of the last elements in the DAF among all replicates has a relative error greater than -0.26 (Figure S10), contrasting with the relative difference of -0.32 observed on real data. This points at a potential additional source of discrepancy between the two methods. To examine whether the differences could be due to selection, we divided the SNPs into annotation categories such as exons, 5’ UTRs, 3’ UTRs, introns, and intergenic regions as mentioned in the polarization above and estimate uSFS for different regions. Since est-sfs only outputs uSFS based on all sites, we re-estimated the uSFS in different categories using the site-specific posterior probabilities from its output (see Methods). However, similar to the results of the polarization section, the differences between the two methods did not show a clear correlation with the expected level of selection intensity in these regions (Figure S12), implying that selection was not the source of the differences in the uSFS from the two methods.

To examine the effect of mutation type on uSFS inference, we oriented mutation types assuming the ancestral state inferred by est-sfs to be true and then compared the relative differences in uSFS estimated by the two methods across mutation types (Figure S13). The relative differences of uSFS of different mutation types showed the same trend as that of the global, that is, the proportion of variants estimated by PolarBEAR in the high DAF category was smaller than that estimated by est-sfs. What’s more, “A->G” and “T->C” together showed a more obvious difference between the two methods; while the other types had much smaller differences, suggesting that the reason for the difference between the two methods is that at least one of the estimations is sensitive to the rates of directional substitutions. When looking at the two methods separately, we found that the estimated uSFS for different mutation types differed only in est-sfs, and the difference was that in “A->G” and “T->C”, a higher proportion of variants were estimated in high DAF categories, but not with PolarBEAR (Figure S14). Unexpectedly, in simulation studies with the presence of only an unequal GC-to-AT content ratio of 3:2 in the ancestral distribution, without involving differences in substitution rates, we found that it can also differentially bias est-sfs estimates for different mutation types (Figure S15). As done for real data, when the polarization results from est-sfs were used to determine the mutation type of the SNPs, the relative differences between the two methods became similar to the differences observed in the real data across mutation types (Figure S16). We note, however, that despite the bias observed for certain mutation types in the simulation, the global uSFS was recovered correctly. Further parameter combinations must be assessed to further characterize the est-sfs bias in the real data.

## Conclusion

PolarBEAR offers a highly accurate polarization method when no outgroup sequence is available, but based on the principle of ARG inference, phased data is necessary.

We show that when the ARG contains information about the ancestral allele state, polarization can be achieved with high accuracy for a subset of variants, depending on their underlying genealogies. Positions with non-informative genealogies do not permit inference with high confidence. Still, we show that posterior probabilities effectively leverage the branch length information to capture the probabilities of ancestral states. A probabilistic approach based on posterior averaging can then be used to infer the unfolded site frequency spectrum accurately. Our result demonstrates the power of ARG-based polarization and uSFS estimation, but also that inference is impacted by errors in the ARG reconstruction. We posit that the accuracy of outgroup-free polarization will further increase with the improvement of ARG inference methods. Our results suggest that pairwise TMRCA inference in combination with a UPGMA tree reconstruction is currently the best option available for the purpose of ancestral states inference and uSFS estimation.

While PolarBEAR could polarize fewer sites than est-sfs on our benchmark dataset, it could analyze SNPs where no outgroup sequence was available. Furthermore, the non-usage of outgroups reduces the issue of substitution model misspecification, which biases uSFS estimates. Therefore, PolarBEAR can serve as a powerful alternative or complementary method for polarizing SNPs.

## Methods

### Simulations

We used msprime (Baumdicker et al. 2022) to simulate ARG and genetic variation data.

#### Default simulation

We simulated one chromosome data with a length of 10M base pair, and with a constant effective population size of 30,000 (60,000 haploids), a mutation rate of 1.25e-8 with the Jukes-Cantor (JC) model (Jukes and Cantor 1969), and the deCODE recombination map (Halldorsson et al. 2019) of human Chr11 starting at position 20004256.

To test the impact of recombination rate, the sites were divided into high and low-recombination rate categories, with the average recombination rate (9.69e-09) as the boundary.

#### Multiple mutations

We generated simulations with a high mutation rate to obtain enough sites to test the impact of multiple mutation events. We used the same ARGs as the default simulations, discarding the original mutations and adding new mutations with a rate of 1e-7 bp^-1^.

#### Demographic scenarios

We simulated two demographic change scenarios, population size increasing from 960 to 46,000 and decreasing from 46,000 to 960, respectively, with a logarithmic change within the last 150,000 generations.

#### Unequal transitions and transversions

As for assessing the effect of multiple mutations, we removed the original--JC-generated--mutations from the ARGs and replaced them with new mutations generated under a Kimura two-parameters substitution model (Kimura 1980) with the transition/transversion rate ratio *κ* = 5.

#### GC-biased root distribution

We simulated ingroup data with population parameters identical to the default simulations but with a simulated sequence length of 50Mb in order to have a sufficient number of sites after classifying them by mutation type. Two outgroup populations with the same size (kept constant) as the ingroup were added, with divergence times at 1.5e6 and 9e5 generations ago, respectively. In non-constant demographic scenarios, changes in population size occur in the ingroup and other parameters are identical to the default simulation scenario. The JC mutation model was used, with equal rates, but a GC-to-AT frequency ratio of 3:2 was used for the root distribution, while equilibrium frequencies remained equal to 0.25.

### ARGs inference methods

Sample sizes with 2, 3, 4, 5, 10, 20, 50, 100 and 200 diploid individuals were used, and ten replicates were performed in each case. Due to the limitation of running time and memory, the maximum sample size for PSMC was 10, while that of other methods was 100. And the sample size 200 was only used for known ARGs.

The PSMC algorithm as implemented in the iSMC program (Barroso et al. 2019) was used with 30 time intervals. The TMRCAs at each SNP positions were computed as the posterior average over all 30 time intervals.

The gamma-SMC (Schweiger and Durbin 2023) program was use with default parameters. The ratio of mutation to recombination rate, was set to 0.78, which is the true ratio in the simulations and also applies to real human data.

The local TMRCAs of the positions with SNPs output by PSMC and gamma-SMC was used to construct a distance matrix describing the haploid relationships. The ‘scipy.cluster.hierarchy.linkage’ function (Virtanen et al. 2020) was used to implement UPGMA clustering to obtain the tree from the distance matrix.

The parameter -t was used in RENT+ (Mirzaei and Wu 2017) to get better branch length estimates. In addition to the control for evaluating the effects of input ancestral states, the major alleles were treated as the ancestral alleles as the input to tsinfer (Kelleher et al. 2019), and if the two alleles each account for 50%, it is treated as missing data. After inferring the tree topology with tsinfer, tsdate (Wohns et al. 2022) was used to estimate branch lengths. For simulated data with demographic changes, the effective population size is calculated using *N*_*e*_ = *θ*/(4*μ*) for the input of tsdate, where the scaled mutation rate *θ* is computed from the data using the Watterson’s estimator.

For PSMC, gamma-SMC and RENT+, the local tree topologies of the positions with SNPs were stored discretely in the tskit tree sequence files. In order to meet tskit’s requirements for node time, the time of non-leaf nodes with time less than 1e-12 were changed to 1e-12; if the time difference between a pair of internal parent and child nodes does not exceed 1e-14, the child node was deleted, which means that polytomies will be generated.

### Polarizing from local genealogies

To determine the most likely ancestral allele, the posterior probabilities of assignments of the states were computed using the empirical Bayesian approach (Yang et al. 1995), and the JC model was used to compute transition probabilities. We performed a joint reconstruction of all ancestral alleles at all inner nodes, as it does not require the averaging of conditional likelihoods at inner nodes, enabling the storage of log probabilities throughout the recursion and avoiding numerical underflow when the tree is large.

Non-informative genealogies, multiple mutations and polytomies are determined based on the detection of mutations in the genealogies, which is achieved through parsimony implemented by the ‘Tree.map_mutations’ function from the tskit package.

### Real data

#### Data Preparation

The phased SNP data and the mask file for chromosome 1 were obtained from the 1000 Genomes Project’s FTP server. Data specific to the African population Mende, Sierra Leone (MSL) containing 87 diploid individuals, was selected for this study. All child samples in trios were excluded.

For the population data, the variants were filtered to include only SNPs that passed all filters. Multiple records of the same positions were collapsed into single multiallelic records. Regions of low confidence or known artefacts were excluded by applying the “StrictMask”.

The MultiZ 20-species alignment file was downloaded from the UCSC Genome Browser. The VCF file including polymorphic and non-polymorphic sites of Human, Orangutan and Macaque was generated using MafFilter (Dutheil et al. 2014). The processed multi-species alignment VCF file was merged with the cleaned population-specific VCF file, keeping only positions present in both sets. Python scripts were used to process the merged VCF file and generate the est-sfs input file.

#### Annotation

The SNPs were annotated using the SnpEff version 5.2c (Cingolani et al. 2012). The reference genome used for annotation was GRCh38, with gene annotations from Ensembl release 99. The classic mode of SnpEff was used.

#### USFS estimation of est-sfs using partial SNPs

First, the posterior probabilities of the major allele as the ancestral state were obtained from the est-sfs output. At the SNP *i*, if the output posterior probability *P*_*output*_ > 0.5, the major allele is the ancestral state, and the posterior probability *P*_*i*_ = *P*_*output*_ ; otherwise, the minor allele is the ancestral state, and the posterior probability *P*_*i*_ = 1 − *P*_*output*_ . Then, for the selected subset of SNPs, uSFS was estimated using the formula used for PolarBEAR in the section *Estimating the unfolded site frequency spectrum*.

## Data Availability

All scripts used in this work are available at https://gitlab.gwdg.de/molsysevol/polarizing_without_outgroup.

## Author contributions

JL: Conceptualization, Methodology, Software, Formal analysis, Investigation, Data curation, Writing - Original Draft, Writing - Review & Editing, Visualization

JYD: Conceptualization, Methodology, Writing - Review & Editing, Supervision

## Acknowledgements

The authors thank Gustavo Barroso, Pengfei Yin, Tal Dagan and Linda Odenthal-Hesse for the helpful suggestions throughout the project. JL was founded by the International Max Planck

Research School for Evolutionary Biology (IMPRS Evol Biol).

## References

Albers PK, McVean G. 2020. Dating genomic variants and shared ancestry in population-scale sequencing data. PLOS Biology 18:e3000586.

Barroso GV, Puzović N, Dutheil JY. 2019. Inference of recombination maps from a single pair of genomes and its application to ancient samples. PLOS Genetics 15:e1008449.

Baumdicker F, Bisschop G, Goldstein D, Gower G, Ragsdale AP, Tsambos G, Zhu S, Eldon B, Ellerman EC, Galloway JG, et al. 2022. Efficient ancestry and mutation simulation with msprime 1.0. Genetics 220:iyab229.

Cingolani P, Platts A, Wang LL, Coon M, Nguyen T, Wang L, Land SJ, Lu X, Ruden DM. 2012. A program for annotating and predicting the effects of single nucleotide polymorphisms, SnpEff. Fly (Austin) 6:80–92.

Collet JM, Nidelet S, Fellous S. 2023. Genetic independence between traits separated by metamorphosis is widespread but varies with biological function. Proceedings of the Royal Society B: Biological Sciences 290:20231784.

Collins TM, Wimberger PH, Naylor GJP. 1994. Compositional Bias, Character-State Bias, and Character-State Reconstruction Using Parsimony. Systematic Biology 43:482–496.

Dutheil JY, Gaillard S, Stukenbrock EH. 2014. MafFilter: a highly flexible and extensible multiple genome alignment files processor. BMC Genomics 15:53.

Eyre-Walker A. 1998. Problems with Parsimony in Sequences of Biased Base Composition. J Mol Evol 47:686–690.

Fang LL, Peede D, Vecchyo DO-D, McTavish EJ, Huerta-Sánchez E. 2024. Leveraging shared ancestral variation to detect local introgression. PLOS Genetics 20:e1010155.

Felsenstein J. 1981. Evolutionary trees from DNA sequences: a maximum likelihood approach. J Mol Evol 17:368– 376.

Glémin S, Arndt PF, Messer PW, Petrov D, Galtier N, Duret L. 2015. Quantification of GC-biased gene conversion in the human genome. Genome Res. 25:1215–1228.

Gutenkunst RN, Hernandez RD, Williamson SH, Bustamante CD. 2009. Inferring the Joint Demographic History of Multiple Populations from Multidimensional SNP Frequency Data. PLOS Genetics 5:e1000695.

Halldorsson BV, Palsson G, Stefansson OA, Jonsson H, Hardarson MT, Eggertsson HP, Gunnarsson B, Oddsson A, Halldorsson GH, Zink F, et al. 2019. Characterizing mutagenic effects of recombination through a sequence-level genetic map. Science 363:eaau1043.

Hernandez RD, Williamson SH, Bustamante CD. 2007. Context Dependence, Ancestral Misidentification, and Spurious Signatures of Natural Selection. Molecular Biology and Evolution 24:1792–1800.

Jukes TH, Cantor CR. 1969. Evolution of Protein Molecules. In: Mammalian Protein Metabolism. Elsevier. p. 21– 132. Available from: https://linkinghub.elsevier.com/retrieve/pii/B9781483232119500097

Keightley PD, Campos JL, Booker TR, Charlesworth B. 2016. Inferring the Frequency Spectrum of Derived Variants to Quantify Adaptive Molecular Evolution in Protein-Coding Genes of Drosophila melanogaster. Genetics 203:975– 984.

Keightley PD, Eyre-Walker A. 2007. Joint Inference of the Distribution of Fitness Effects of Deleterious Mutations and Population Demography Based on Nucleotide Polymorphism Frequencies. Genetics 177:2251–2261.

Keightley PD, Jackson BC. 2018. Inferring the Probability of the Derived vs. the Ancestral Allelic State at a Polymorphic Site. Genetics 209:897.

Kelleher J, Wong Y, Wohns AW, Fadil C, Albers PK, McVean G. 2019. Inferring whole-genome histories in large population datasets. Nat Genet 51:1330–1338.

Kimura M. 1980. A simple method for estimating evolutionary rates of base substitutions through comparative studies of nucleotide sequences. J Mol Evol 16:111–120.

Koenig D, Hagmann J, Li R, Bemm F, Slotte T, Neuffer B, Wright SI, Weigel D. 2019. Long-term balancing selection drives evolution of immunity genes in Capsella.Przeworski M, Baldwin IT, Gao Z, editors. eLife 8:e43606.

Langley CH, Stevens K, Cardeno C, Lee YCG, Schrider DR, Pool JE, Langley SA, Suarez C, Corbett-Detig RB, Kolaczkowski B, et al. 2012. Genomic Variation in Natural Populations of Drosophila melanogaster. Genetics 192:533.

Li H, Durbin R. 2011. Inference of Human Population History From Whole Genome Sequence of A Single Individual. Nature 475:493.

Marchi N, Excoffier L. 2020. Gene flow as a simple cause for an excess of high-frequency-derived alleles. Evolutionary Applications 13:2254–2263.

Mirzaei S, Wu Y. 2017. RENT+: an improved method for inferring local genealogical trees from haplotypes with recombination. Bioinformatics 33:1021–1030.

Pivirotto AM, Platt A, Patel R, Kumar S, Hey J. 2024. Analyses of allele age and fitness impact reveal human beneficial alleles to be older than neutral controls. eLife [Internet] 13. Available from: https://elifesciences.org/reviewed-preprints/93258

Sabeti PC, Varilly P, Fry B, Lohmueller J, Hostetter E, Cotsapas C, Xie X, Byrne EH, McCarroll SA, Gaudet R, et al. 2007. Genome-wide detection and characterization of positive selection in human populations. Nature 449:913.

Schaefer NK, Shapiro B, Green RE. 2021. An ancestral recombination graph of human, Neanderthal, and Denisovan genomes. Science Advances 7:eabc0776.

Schneider A, Charlesworth B, Eyre-Walker A, Keightley PD. 2011. A Method for Inferring the Rate of Occurrence and Fitness Effects of Advantageous Mutations. Genetics 189:1427–1437.

Schweiger R, Durbin R. 2023. Ultrafast genome-wide inference of pairwise coalescence times. Genome Research 33:1023.

Sendrowski J, Bataillon T. 2024. fastDFE: Fast and Flexible Inference of the Distribution of Fitness Effects. Molecular Biology and Evolution 41:msae070.

Speidel L, Forest M, Shi S, Myers SR. 2019. A method for genome-wide genealogy estimation for thousands of samples. Nat Genet 51:1321–1329.

Stern AJ, Wilton PR, Nielsen R. 2019. An approximate full-likelihood method for inferring selection and allele frequency trajectories from DNA sequence data. PLOS Genetics 15:e1008384.

Virtanen P, Gommers R, Oliphant TE, Haberland M, Reddy T, Cournapeau D, Burovski E, Peterson P, Weckesser W, Bright J, et al. 2020. SciPy 1.0: fundamental algorithms for scientific computing in Python. Nat Methods 17:261– 272.

Voight BF, Kudaravalli S, Wen X, Pritchard JK. 2006. A Map of Recent Positive Selection in the Human Genome. PLoS Biology 4:e72.

Wohns AW, Wong Y, Jeffery B, Akbari A, Mallick S, Pinhasi R, Patterson N, Reich D, Kelleher J, McVean G. 2022. A unified genealogy of modern and ancient genomes. Science 375:eabi8264.

Yang Z, Kumar S, Nei M. 1995. A New Method of Inference of Ancestral Nucleotide and Amino Acid Sequences. Genetics 141:1641.

Zeng K, Fu Y-X, Shi S, Wu C-I. 2006. Statistical Tests for Detecting Positive Selection by Utilizing High-Frequency Variants. Genetics 174:1431.

