## Supplementary Information for "Polarizing SNPs without outgroup"

### Supp. Text 1: Branch lengths influence the posterior probabilities of ancestral alleles in non-informative genealogies

If we assume a non-informative genealogy:

1. two branches are starting from the root node connecting to the nodes $N_1$ and $N_2$ with branch lengths of $L_1$ and $L_2$;

2. the genotype of all samples under $N_1$ is $G_1$, and the genotype of all samples under $N_2$ is $G_2$;

3. the rate of changing from $G_1$ to $G_2$ is equal to the rate of changing from $G_2$ to $G_1$, denoted as $\theta$.

The probability of a mutation of a certain type is denoted as

$$\mathrm{Mut}(BranchLength, \theta) = \frac{1}{3} \times \left(1 - e^{-BranchLength \times \theta}\right)$$

The probability of mutation doesn't occur is denoted as

$$\mathrm{NoMut}(BranchLength, \theta) = e^{-BranchLength \times \theta}$$

Because the mutation rate is low, the likelihood that the genotypes at two nodes are the observed genotypes in their descendant samples is close to 1. Under the infinite site model, the mutation must have occurred in one of the two branches from the root, so there couldn't be any mutation under the $N_1$ and $N_2$ nodes. So, the observed genotype $G_1$ and $G_2$ at $N_1$ and $N_2$ can be seen as given conditions. Instead of polarizing along the entire tree, it can be viewed as polarizing only along a tree with a root node and two "ancient samples" ($N_1$ and $N_2$) with different branch lengths.

So, the probability of $G_1$ as genotype of the root is:

$$\mathrm{P1} \approx \mathrm{NoMut}(L_1, \theta) \times \mathrm{Mut}(L_2, \theta)$$

the probability of $G_2$ as genotype of the root is:

$$\mathrm{P2} \approx \mathrm{Mut}(L_1, \theta) \times \mathrm{NoMut}(L_2, \theta)$$

Expand $\mathrm{Mut}(BranchLength, \theta)$ and $\mathrm{NoMut}(BranchLength, \theta)$:

$$\mathrm{P1} = \mathrm{NoMut}(L_1, \theta) \times \mathrm{Mut}(L_2, \theta)$$

$$= e^{-L_1 \times \theta} \times 1/3 \times (1 - e^{-L_2 \times \theta})$$

$$= 1/3 \times (e^{-L_1 \times \theta} - e^{-(L_1 + L_2) \times \theta})$$

similarly,

$$\mathrm{P2} = \mathrm{Mut}(L_1, \theta) \times \mathrm{NoMut}(L_2, \theta) = 1/3 \times (e^{-L_2 \times \theta} - e^{-(L_1 + L_2) \times \theta})$$

So, the ratio of $\mathrm{P1}$ to $\mathrm{P2}$ is,

$$\frac{\mathrm{P1}}{P2} = \frac{e^{-L_1 \times \theta} - e^{-(L_1 + L_2) \times \theta}}{e^{-L_2 \times \theta} - e^{-(L_1 + L_2) \times \theta}}$$

Divided both $\mathrm{P1}$ and $\mathrm{P2}$ by $e^{-(L_1 + L_2) \times \theta}$,

$$\frac{\mathrm{P1}}{P2} = \frac{e^{(-L_1 + L_1 + L_2) \times \theta} - 1}{e^{(-L_2 + L_1 + L_2) \times \theta} - 1} = \frac{e^{L_2 \times \theta} - 1}{e^{L_1 \times \theta} - 1}$$

When $L_2 \times \theta \to 0$ *and* $L_1 \times \theta \to 0$:

$$e^{L_2 \times \theta} \cong L_2 \times \theta + 1$$

$$e^{L_1 \times \theta} \cong L_1 \times \theta + 1$$

so,

$$\frac{\mathrm{P1}}{P2} \cong \frac{L_2 \times \theta}{L_1 \times \theta} = \frac{L_2}{L_1}$$

If $L_2 > L_1$, $G_1$ is the ancestral state, with posterior probability $P \cong L_1/(L_1 + L_2)$.

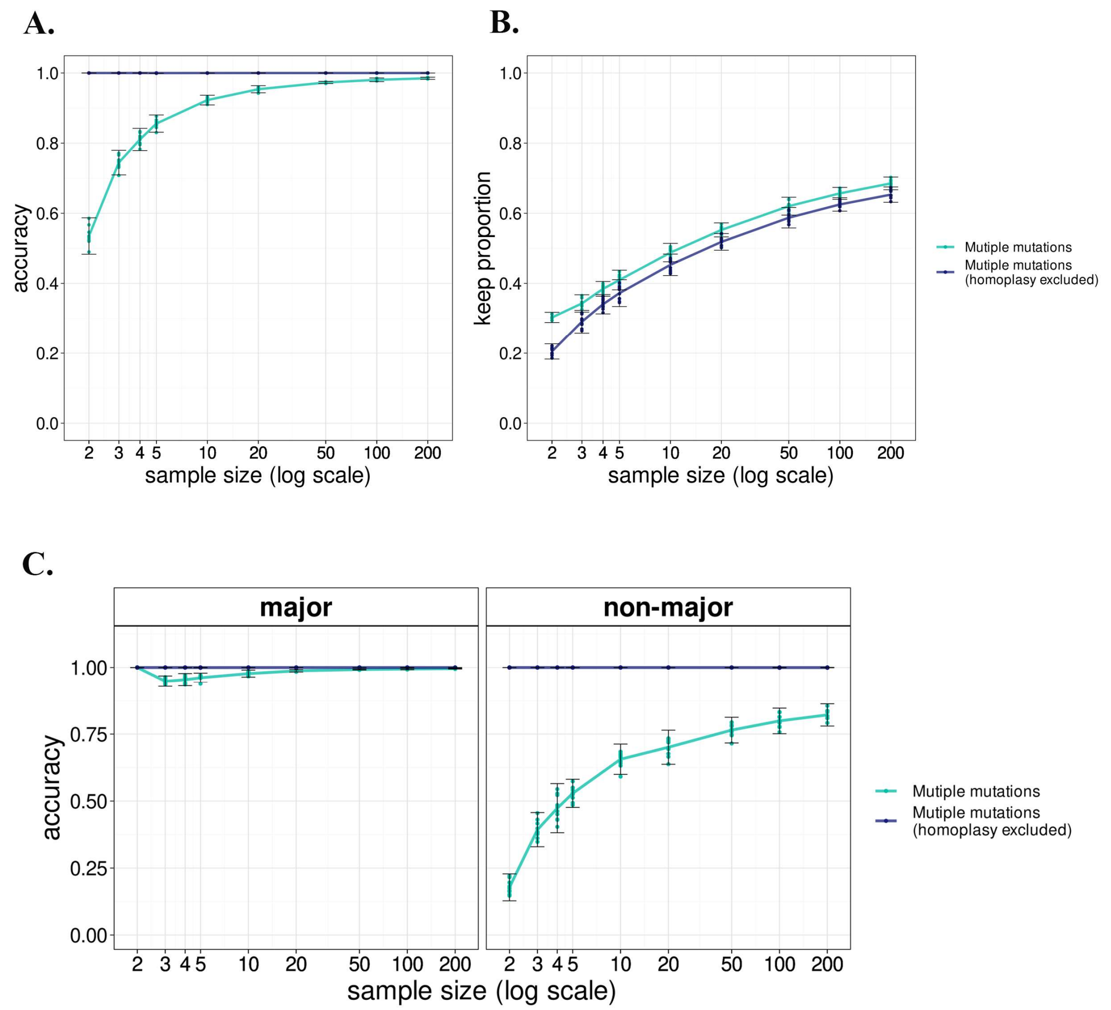


**Figure S1. Polarizing performance of PolarBEAR for sites with multiple mutations events with simulated ARG**. Total accuracy (A), proportion of kept SNPs (B) and accuracy of among the SNPs classified by their ancestral alleles as major and minor (C) for different sample sizes, with and without excluding the homoplasy sites. The points show 10 replicates in each group, and the lines represent their means.

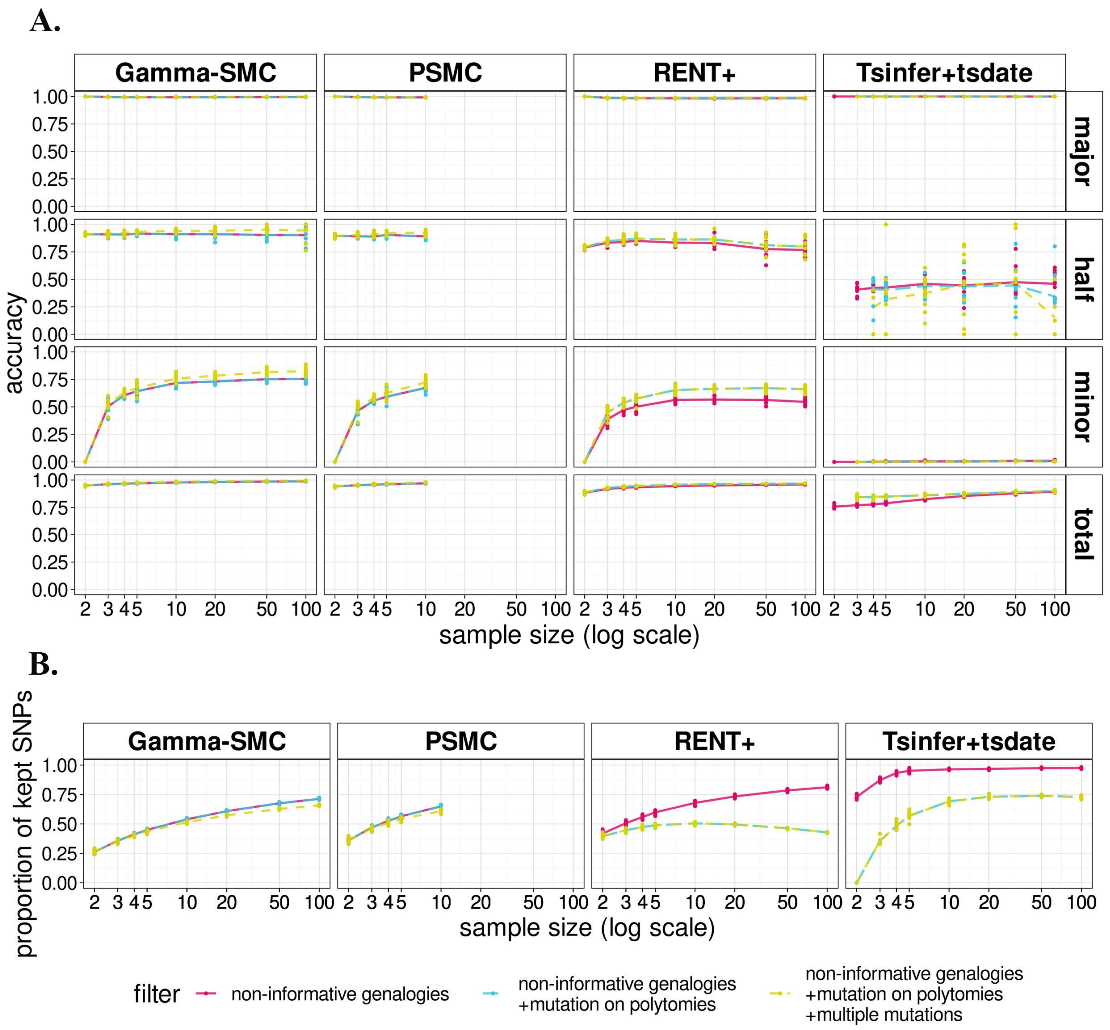


**Figure S2. Effects of different filtering schemes on the performance of PolarBEAR with ARGs inferred by different methods.** Filter 1: only non-informative genealogies are filtered. Filter 2: Filter 1 + sites where mutations occur at polytomies are filtered. Filter 3: Filter 2 + sites that require more than one mutation (as inferred by maximum parsimony reconstruction) to explain the distribution of the genotypes on leaves are filtered, represented by different colors. Accuracy comparison for SNPs classified by their ancestral allele frequencies (major, equal, or minor, A), and proportion of kept SNPs (B) for different sample sizes. The points show 10 replicates in each group, and the lines represent their means.

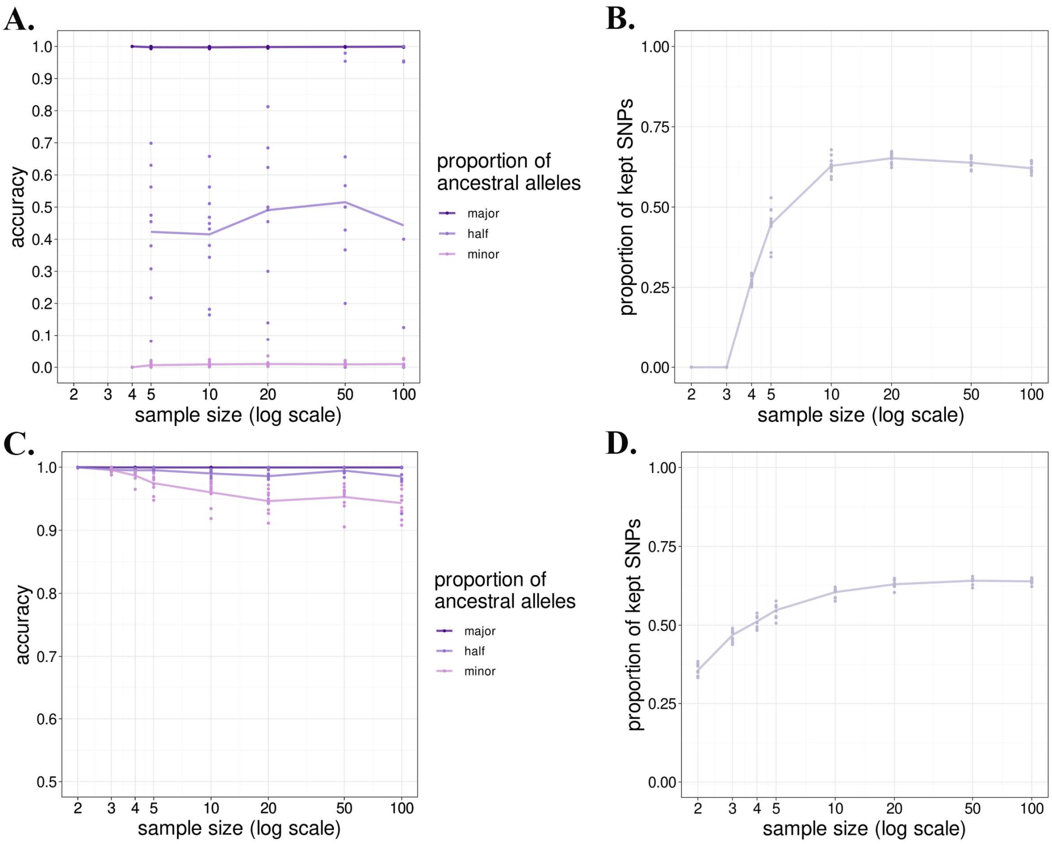


**Figure S3. Polarizing performance of PolarBEAR with tsinfer+tsdate using the first round of polarization results with all filters (A and B) and true ancestral alleles (C and D) as input**. Comparison of polarization accuracy of SNPs classified by the frequency of their ancestral alleles (major, equal, or minor) shown in different colors (A and C) and proportion of analyzed SNPs after filtering (B and D) for different sample sizes.

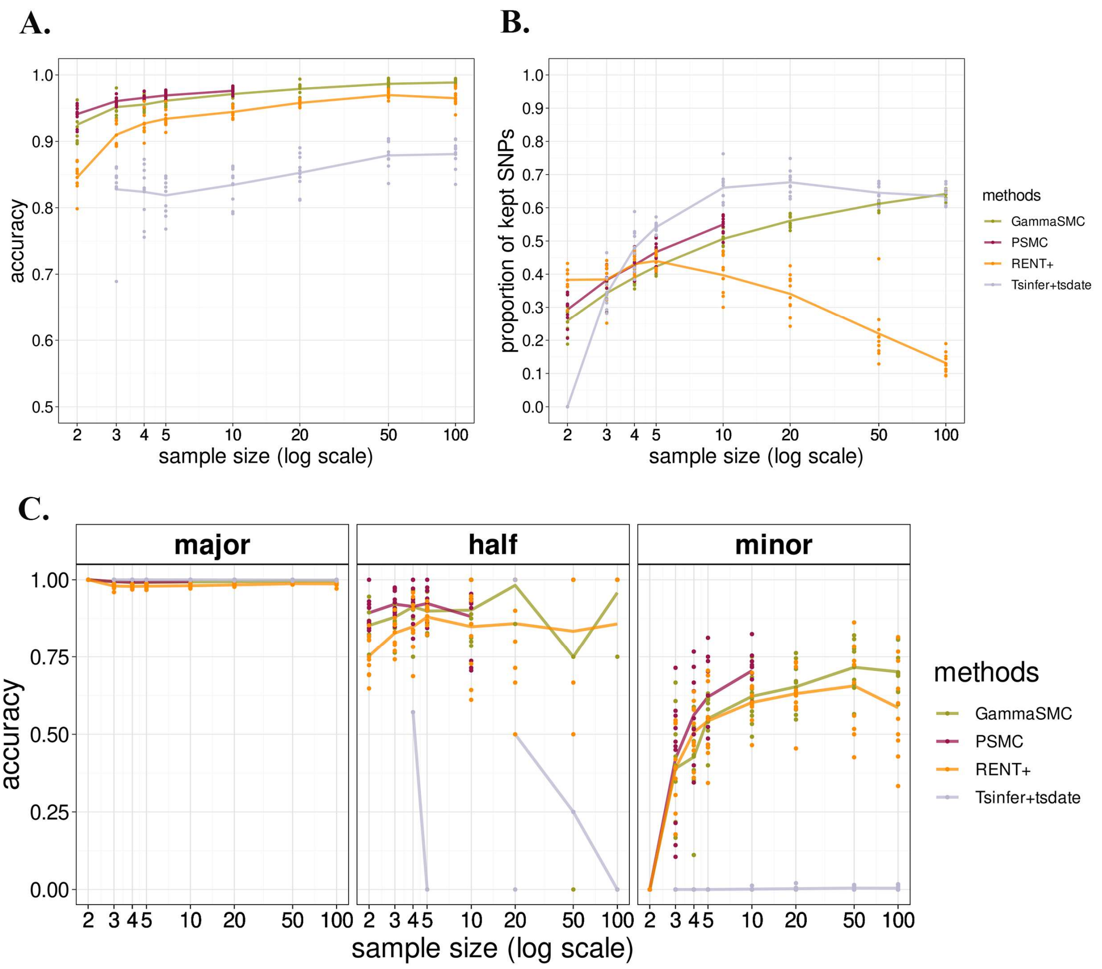


**Figure S4. Polarizing performance of PolarBEAR with inferred ARGs for a declining population size scenario**. Legend as in Figure 2.

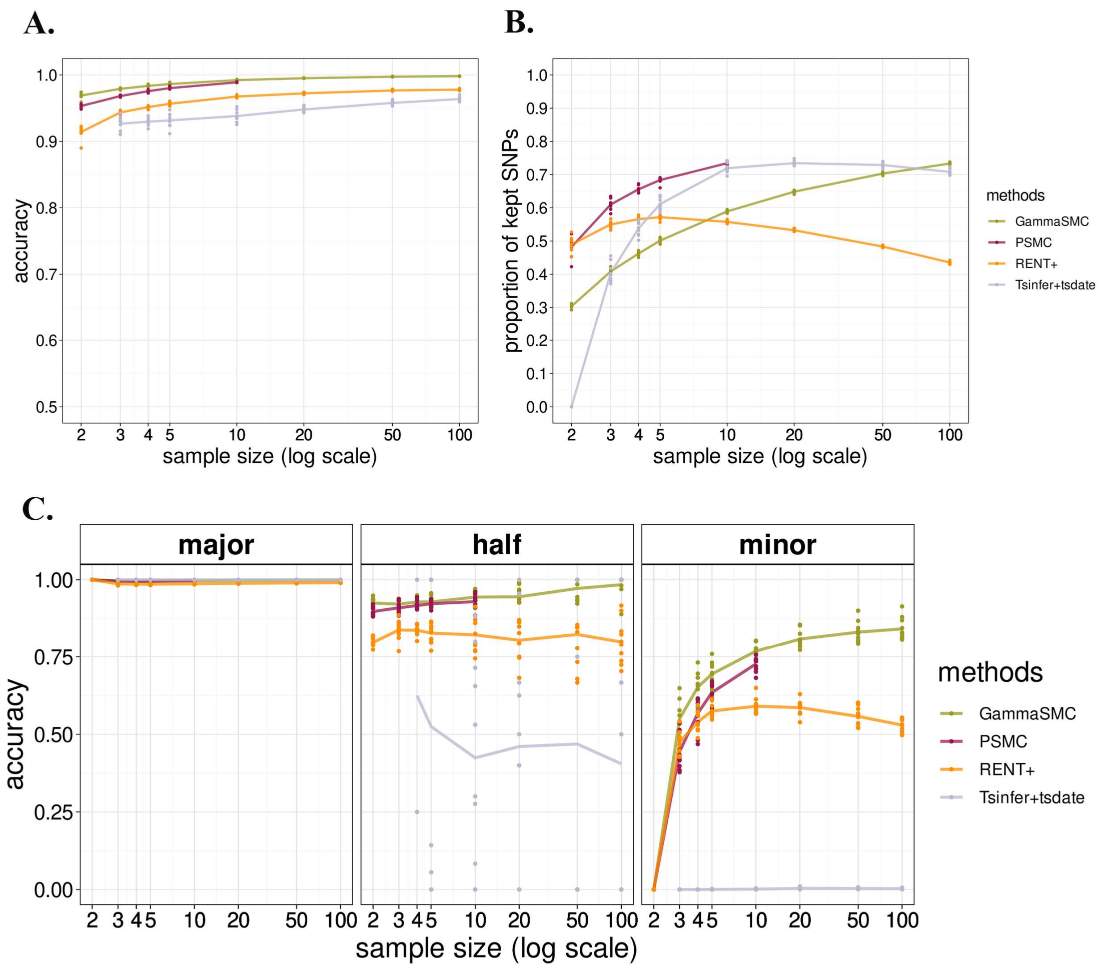


**Figure S5. Polarizing performance of PolarBEAR with inferred ARGs for an increasing population size scenario**. Legend as in Figure 2.

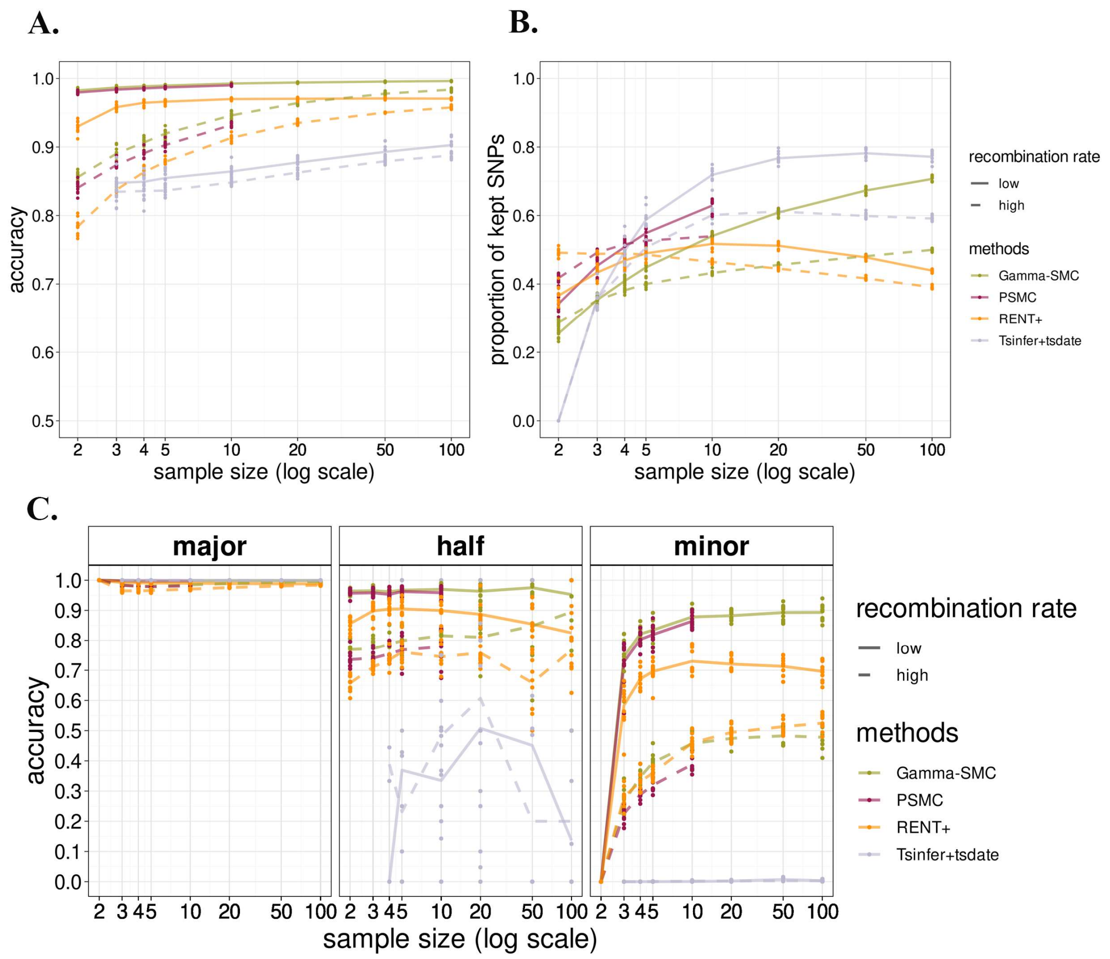


**Figure S6. Effects of high and low recombination rates (shown in dashed and solid line) on the performance of PolarBEAR with ARGs inferred by gamma-SMC, PSMC, RENT+, and tsinfer+tsdate (shown in different colors)**. Comparison of total accuracy (A), proportion of analyzed SNPs after filtering (B) and accuracy for SNPs classified by their ancestral allele frequency (major, equal, or minor, C) for different sample sizes. The points show 10 replicates in each group, and the lines represent their means.

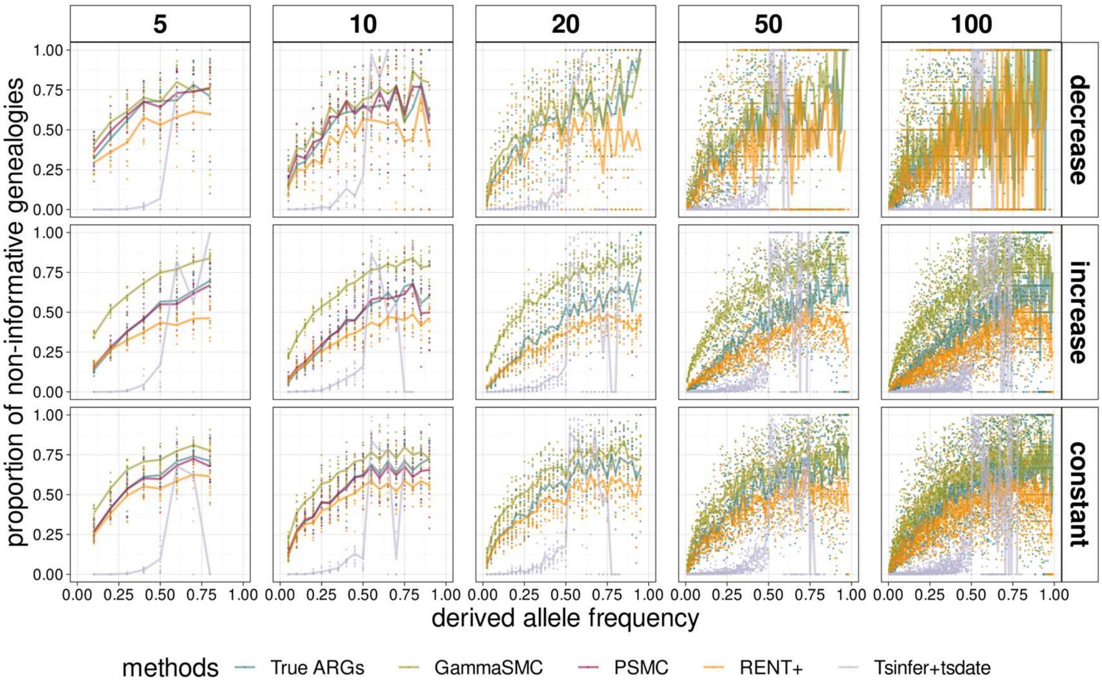


**Figure S7. Proportion of SNPs with non-informative genealogies as a function of the derived allele frequency**. Ancestral alleles were inferred by PolarBEAR with gamma-SMC, PSMC, RENT+, and tsinfer+tsdate (in different colors), for different sample sizes (in different columns) and under different demographic scenarios (in different rows).

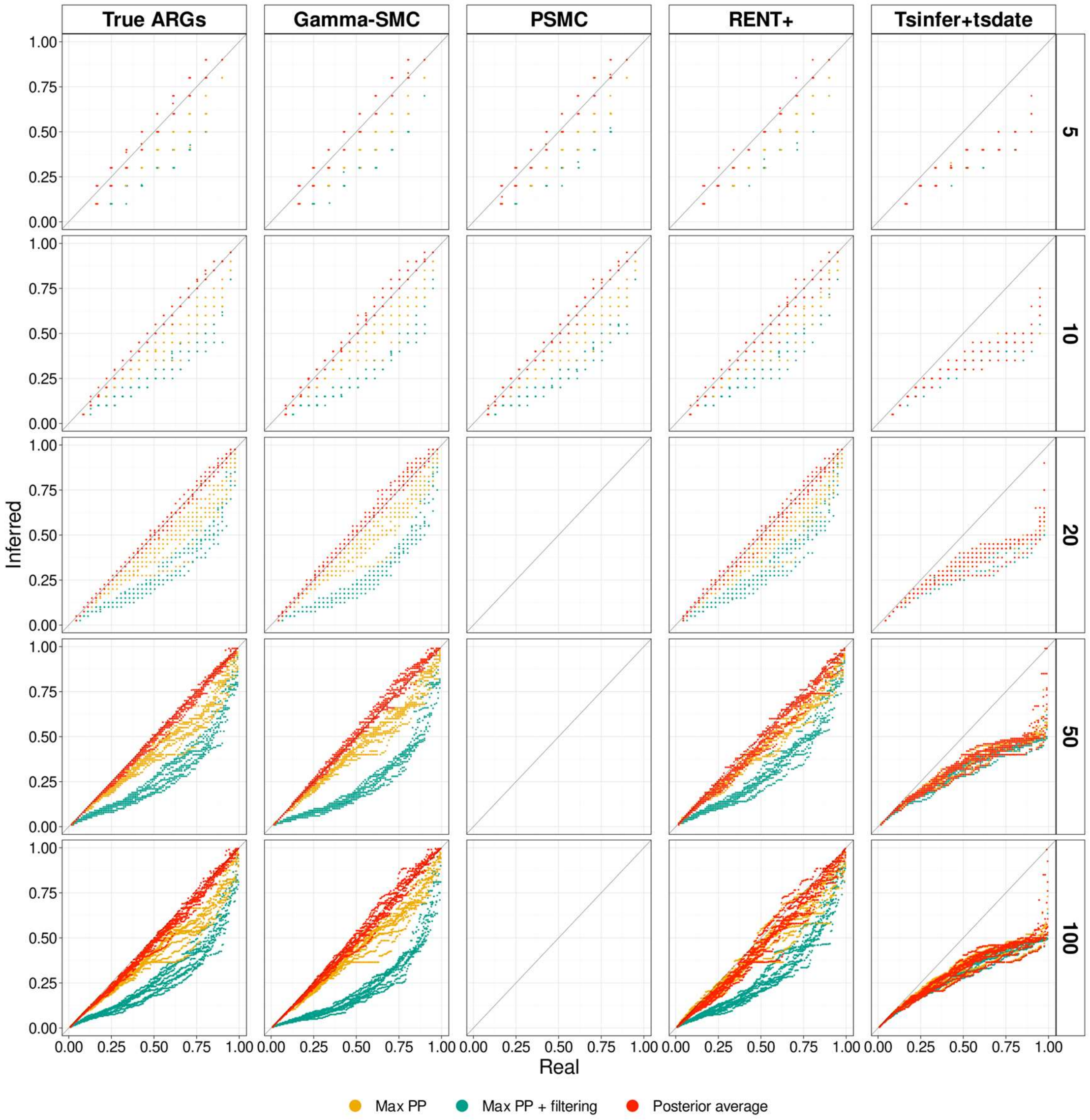


**Figure S8. uSFS inference with different ARG reconstruction methods under a decreasing population size scenario**. Legend as in Figure 3.

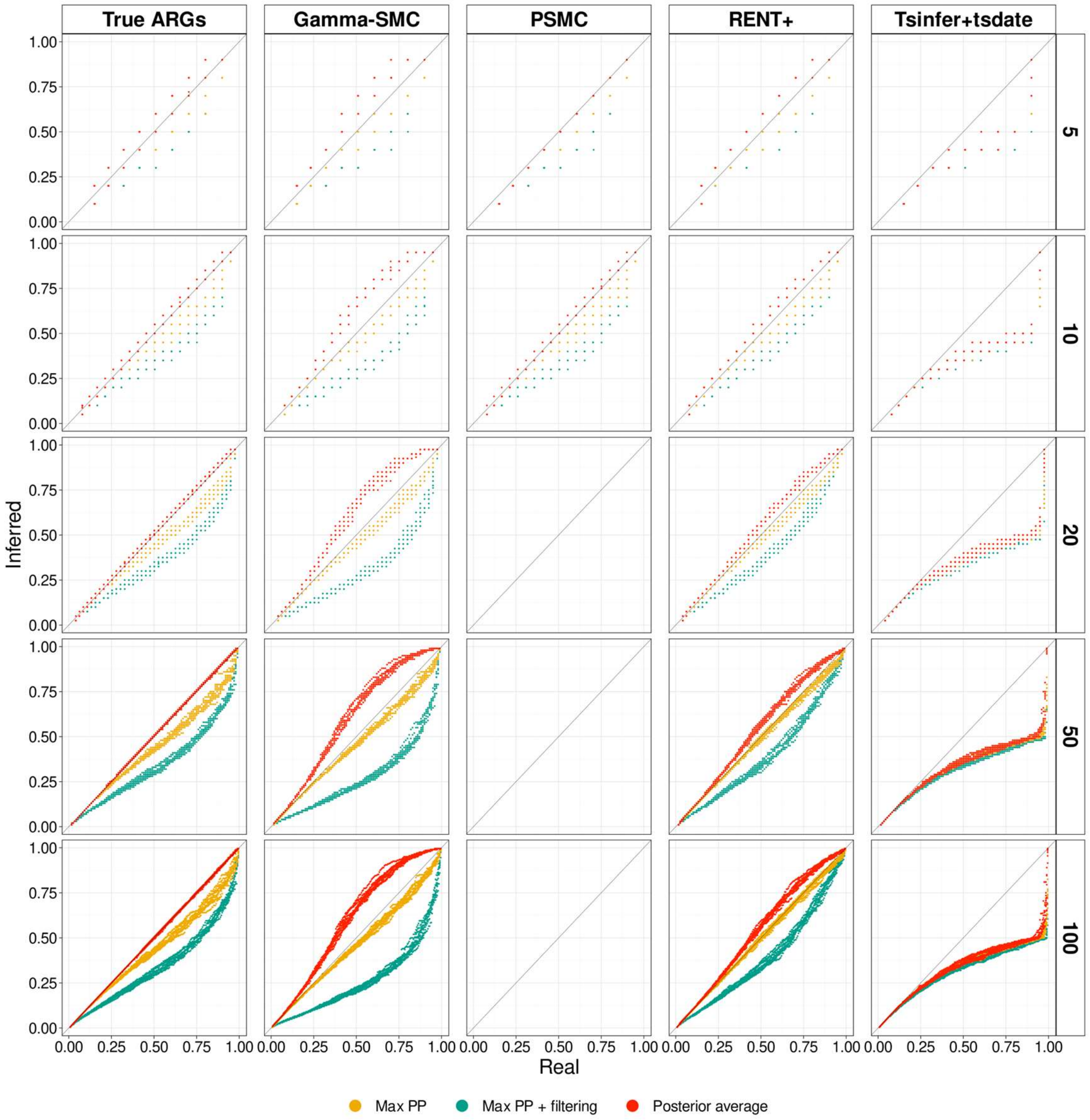


**Figure S9. uSFS inference with different ARG reconstruction methods under an increasing population size scenario**. Legend as in Figure 3.

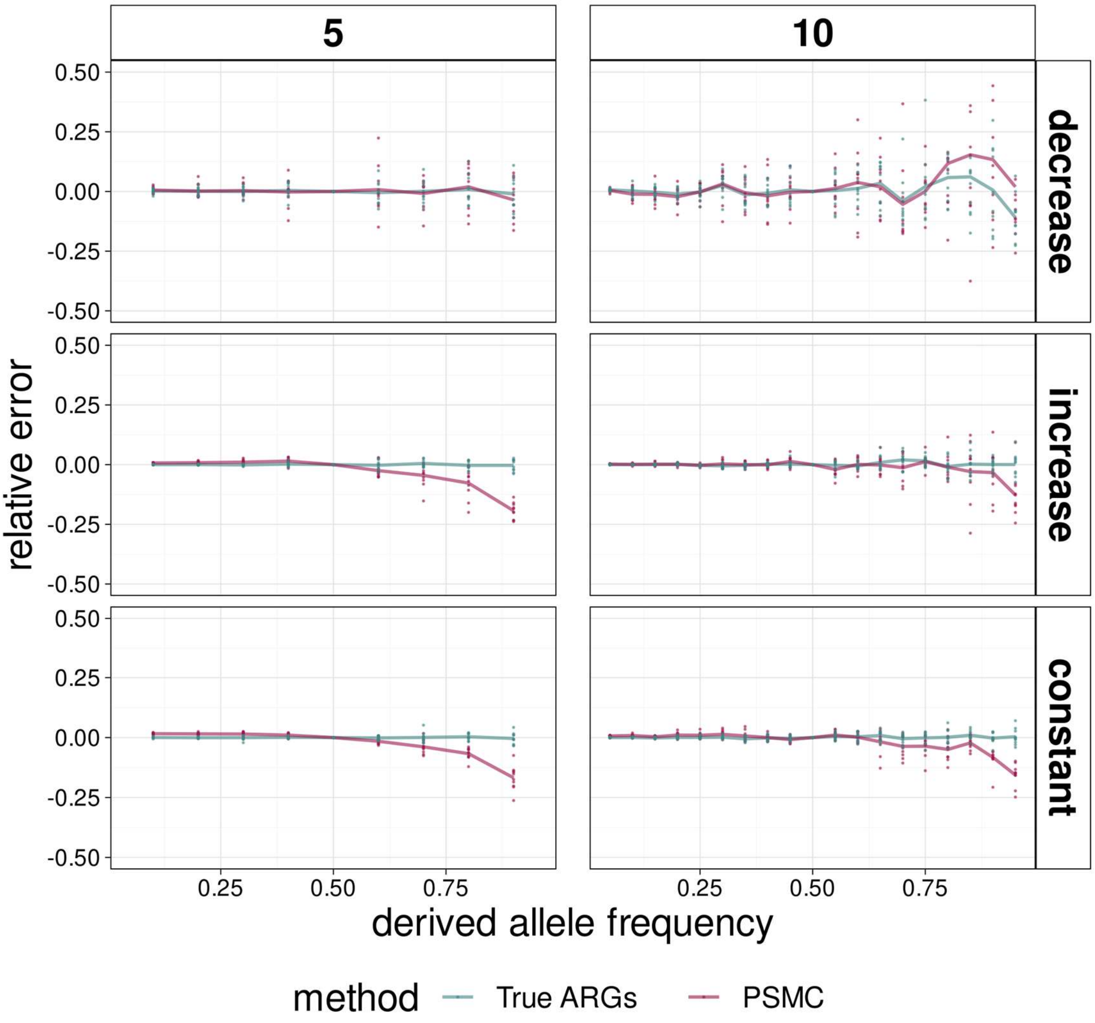


**Figure S10. Relative error of uSFS inference from simulations**. uSFS estimated by PolarBEAR with true ARGs (blue) and ARGs from PSMC (pink), under different demographic scenarios (rows) and sample sizes columns). The points show 10 replicates in each group, and the lines represent their means. Some of the points with large variability are not shown because of the y-axis limits, especially in decreasing population sizes.

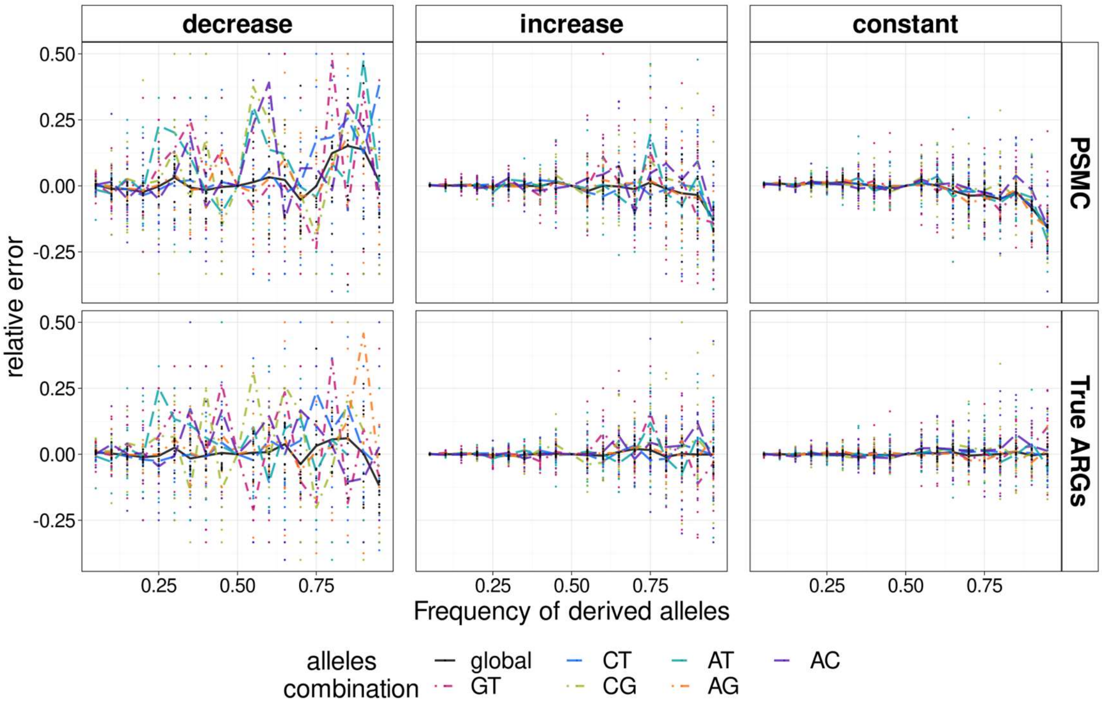


**Figure S11. Relative error of uSFS estimated from simulations under distinct transition and transversion rates.** Colors show different allele combinations. Ancestral alleles estimated by PolarBEAR with true ARGs and ARGs from PSMC (rows), under simulations with different demographic scenarios (columns). Ten replicates were simulated in each case using Kimura's two-parameter model with $\kappa = 5$. The points show 10 replicates in each group, and the lines represent their means. Some of the points with large variability are not shown because of the y-axis limits, especially in decreasing population sizes.

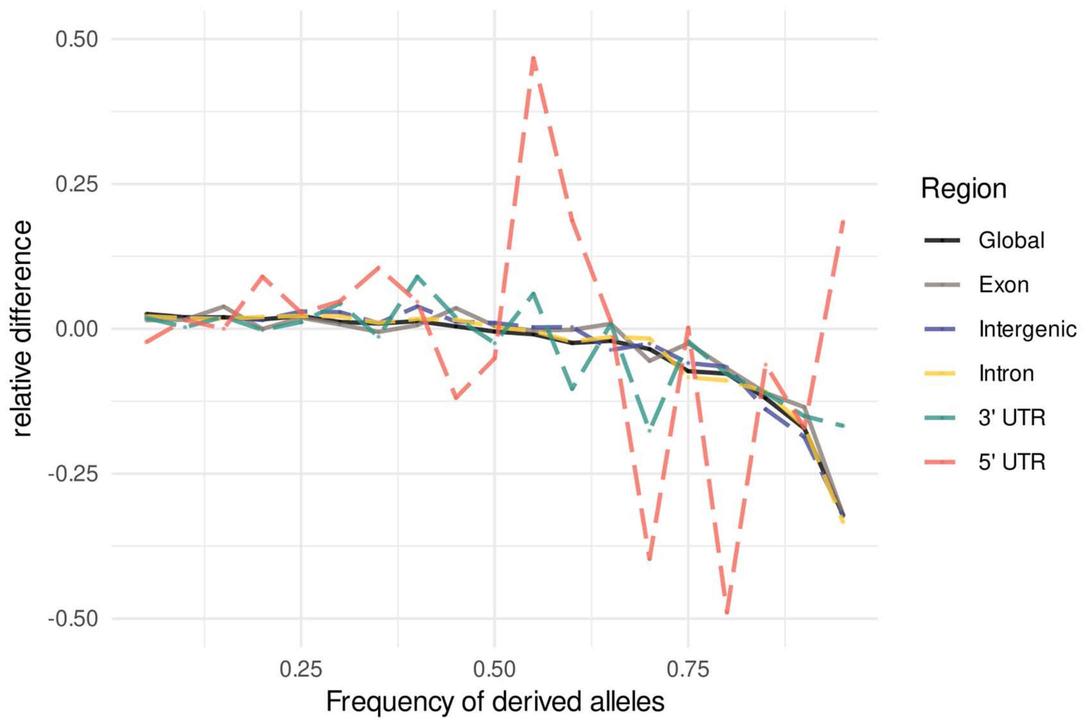


**Figure S12. Relative difference between uSFS estimated by PolarBEAR and est-sfs in different genomic regions**. The y-axis displays $(PolarBEAR - est\_sfs)\big/_{est\_sfs}$, with SNPs annotated to be in exon, intron, 5’ UTR, 3’UTR and intergenic regions, and the full chromosome (in different colors). Data from human chromosome 1.

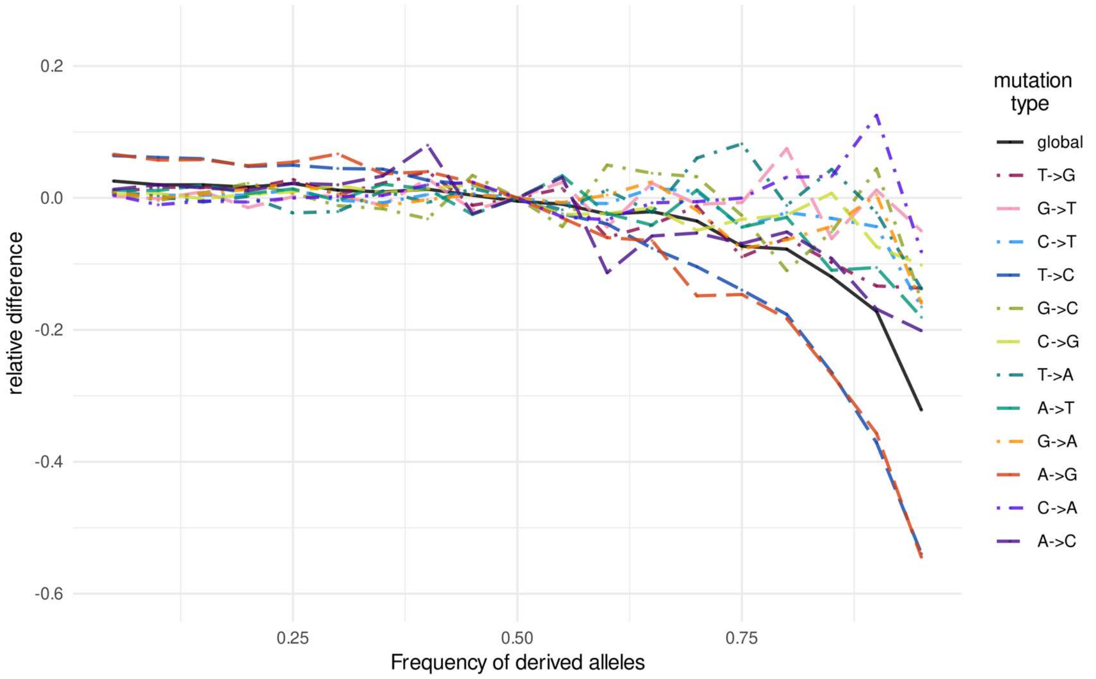


**Figure S13. Relative difference between uSFS estimated by PolarBEAR and est-sfs, for distinct mutation types**. The y-axis displays $(PolarBEAR - est\_sfs)/est\_sfs$, with SNPs with different mutation types (in different colors) oriented by polarized states from est-sfs. Data from human chromosome 1.

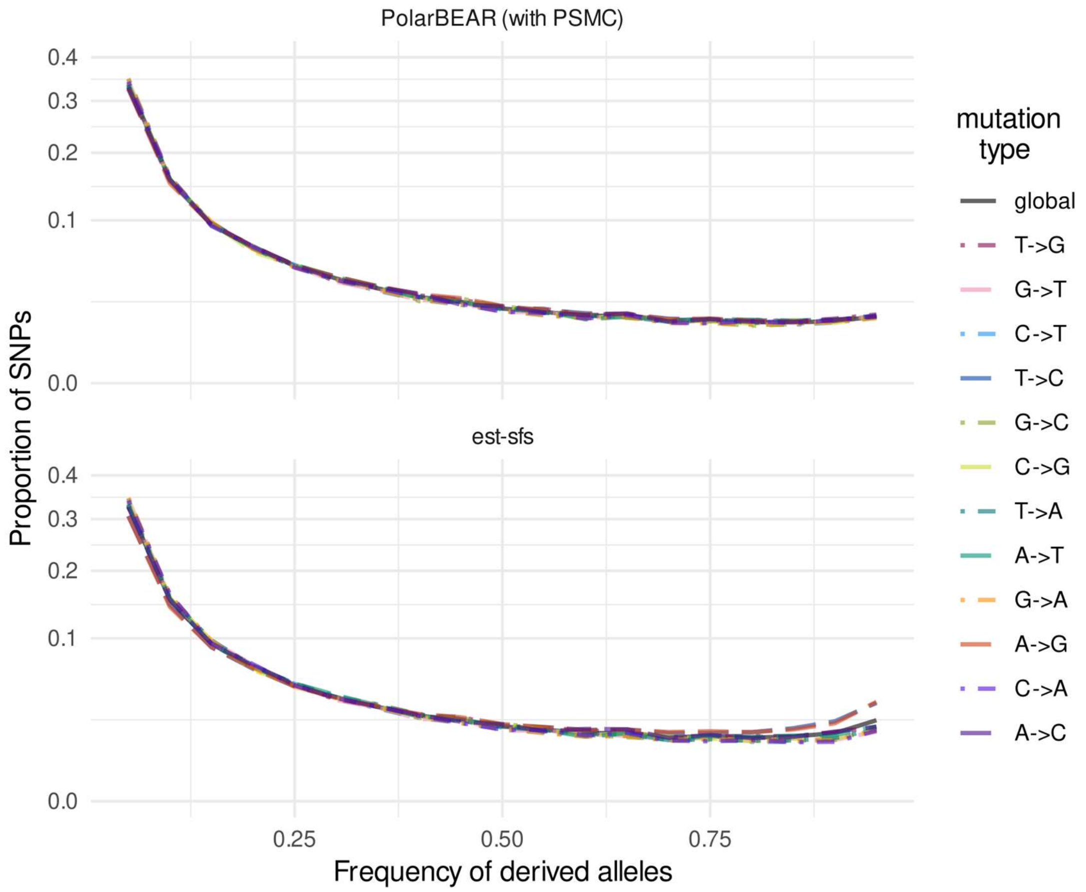


**Figure S14. Estimated uSFS for different mutation types.** Unfolded SFS estimated by PolarBEAR with PSMC (top) and est-sfs (bottom), SNPs with different mutation types (in different colors) oriented by polarized states from est-sfs. The y-axis, the proportion of SNPs, is scaled using a square root transformation. Data from human chromosome 1.

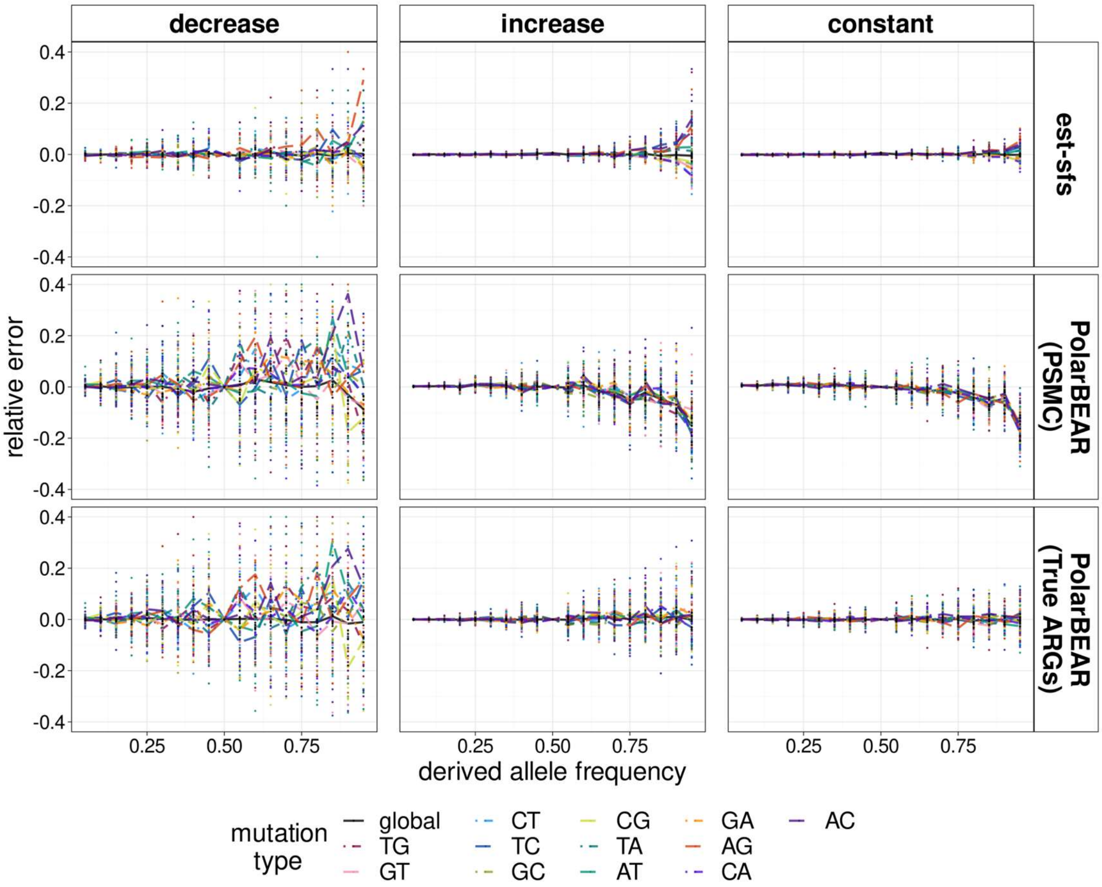


**Figure S15. Relative error of uSFS estimated from simulations under unequal GC content.** Legend as in figure S11, with the addition row of relative error of uSFS estimated by est-sfs. Simulations with unequal GC content (60%) in the ancestral distribution.

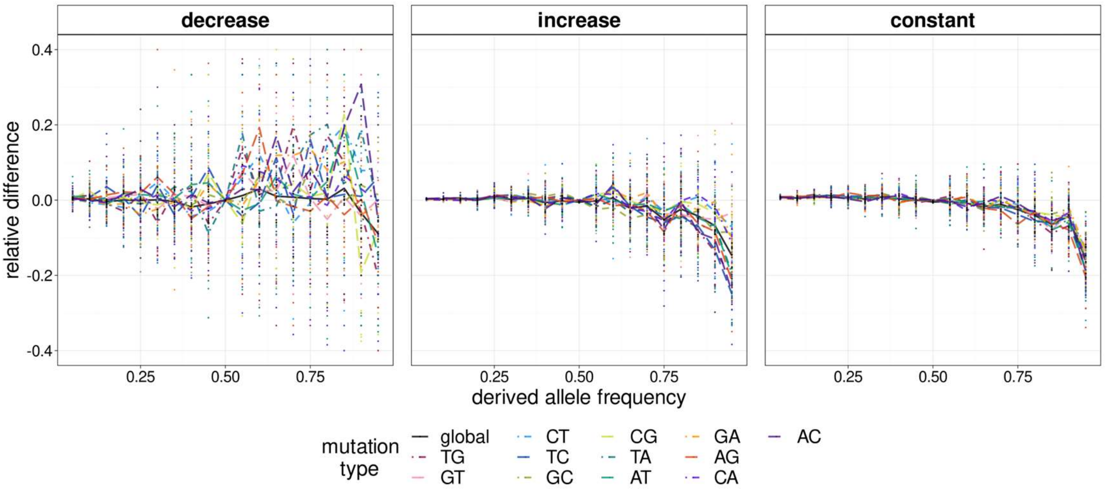


**Figure S16. Relative differences between uSFS estimated by PolarBEAR and est-sfs under unequal GC content.** Legend as in Figure S13. Data simulated with unequal GC content (60%) in the ancestral distribution and under distinct demographic scenarios.
